# Molecular map of GNAO1-related disease phenotypes and reactions to therapy

**DOI:** 10.1101/232058

**Authors:** Ivana Mihalek, Jeff L. Waugh, Meredith Park, Saima Kayani, Annapurna Poduri, Olaf Bodamer

**Affiliations:** Department of Molecular Medicine and Biotechnology, Faculty of Medicine, University of Rijeka, Rijeka, Croatia; Department of Pediatrics, University of Texas Southwestern Medical Center, Dallas, TX; Department of Neurology, Boston Children’s Hospital, Harvard Medical School, Boston, MA; Epilepsy Genetics Program, Department of Neurology, Boston Children’s Hospital, Boston, MA; currently: University of North Carolina at Chapel Hill School of Medicine; F.M. Kirby Neurobiology Center, Boston Children’s Hospital, Boston, MA; Broad Institute of MIT and Harvard University, Cambridge, MA; Division of Genetics and Genomics, Boston Children’s Hospital, Harvard Medical School, Boston, MA

## Abstract

The GNAO1 gene codes for the most commonly expressed G*α* protein in the central nervous system. Pathogenic GNAO1 variants result in early-onset neurological phenotypes, sometimes with distinct epilepsy or movement disorder, and sometimes with both manifestations in the same patient. The existing extensive knowledge about G-protein coupled receptor (GPCR) signaling provides the input needed to describe quantitatively how mutations modify the GPCR signal. This in turn allows rational interpretation of distinct phenotypes arising from mutations in GNAO1. In this work we outline a model that enables understanding of clinical phenotypes at a molecular level. The mutations affecting the catalytic pocket of GNAO1, we show, result in the improper withdrawal of the signal, and give rise to epileptic phenotypes (EPs). The converse is not true - some pure EPs are caused by mutations with no obvious impact on catalysis. Mutations close to the interface with GNAO1’s downstream effector block the signal propagation in that direction, and manifest as a movement disorder phenotype without epilepsy. Quantifying the reported reaction to therapy highlights the tendency of the latter group to be unresponsive to the therapies currently in use. We argue, however, that the majority of clinically described mutations can impact several aspects of GNAO1 function at once, resulting in the continuum of phenotypes observed in patients. The reasoning based on GNAO1 signaling model provides a precision medicine paradigm to aid clinicians in selecting effective categories of medication, and in addition, can suggest pragmatic targets for future therapies.

## Introduction

From the reports [1, 2, 3, 4, 5], and early compilations [6, 7, 8, 9, 10], of disease-associated variants in GNAO1 gene, a picture emerges of a source of complex phenotypes, with equally complex and sometimes contradictory responses to treatment. GNAO1 (guanine nucleotide binding protein, subunit *α*, type “o”) is the most common type of G*α* in the central nervous system (CNS). It functions at the core of multiple neurotransmitter pathways, and is expressed in multiple types of neurons [11]. In particular, GNAO1 expresses profusely in the striatum, [12, 13], Table S1, region of the brain responsible for the smooth transition between movement sequences [14]. Unsurprisingly then, GNAO1 pathogenic variants cause a spectrum of neurological phenotypes, roughly falling into two categories: movement disorders (MD) or epilepsy-related (E) phenotypes. While most patients manifest symptoms from both categories, some patients have exclusively MD or E manifestations. The most frequent MD phenotype is chorea, though the spectrum can include or be limited to myoclonus or dystonia. Epilepsy related to GNAO1 is typically early-onset, often with status epilepticus and/or seizures that are refractory to multiple anti-seizure medication regimens. The epilepsy-dominant patients typically have global developmental delay, often accompanied by severe hypotonia, and manifest stereotypies and difficulty with motor coordination. The fact that the two patterns may completely segregate points to two distinct classes of molecular abnormality underlying GNAO1-related symptoms. At the same time, the movement and epilepsy phenotypes do overlap in the majority of cases, suggesting that a single mutation can impact multiple aspects of GNAO1 function, eluding a straightforward gain-and loss-of-function classification [15].

Prior reports on GNAO1 therapy and its effectiveness have not systematically characterized medication responses on both the epilepsy and movement disorders axes. However, if we consider drugs in terms of their main targets, roughly two classes of disease variants emerge: one responsive to modulation of GABA and dopamine receptors, and the other refractory to all reported therapy. In large part, the first class overlaps with the variants close to the catalytic pocket of GNAO1 [10], while the second overlaps with those close to the GNAO1-effector interface. The overlap is incomplete, and the therapeutic response much more complex than this schematic classification. The challenge thus lies in characterizing the molecular lesions in a way that reconciles the seemingly contradictory treatment requirements and predicts medication efficacy, potentially reducing side effects and time to benefit.

Here we show that even though the concrete aspects of GNAO1-related phenotype may depend on the individual genetic background and development history, the main features of the genotype-phenotype correlation can be rationalized by limiting our focus on the core loop of the G protein signaling pathway, and the way in which disease-related mutations modify its signal.

## Background: Molecular players in GNAO1 signaling pathway

We describe a model that centers on the three obligate components of the GPCR signaling pathway [16], Fig. S4 and Movie S1: a GTPase, a guanine nucleotide exchange factor (GEF), and a GTPase-activating protein (GAP). It is a particular feature of GPCR signaling, in contrast to the signaling through small GTPases, that the GTPase, G*α*, forms a complex with G*βγ* [17]. This complex dissociates upon the interaction with activated GPCR, creating a branching, or bipartite, signal [18], with G*α* and G*βγ* independently influencing their downstream effectors.

### GTPase: GNAO1

GNAO1 is severalfold more abundant than any other G*α* protein in the CNS [11]. Neurons and neuroendocrine cells are the primary site of expression of the gene [19, 13]. Known G*βγ* partners of GNAO1 are encoded by GNB1 (as G*β*) and GNG2 (as G*γ*) [20], though the existence of other partners is likely. As is the case with GNAO1, GNB1 and GNG2 are preferentially expressed in the brain [21]. Of the five genes coding for G*β* [22], GNB1 is the only one that has been related to autosomal dominant neurodevelopmental disease phenotypes, typically global developmental delays and hypotonia, sometimes accompanied by epilepsy and/or dystonia [23, 24]. Known disease-related GNB1 mutations are located almost exclusively at the interface with G*α* [24], Movie S1.

### GEF: G-protein coupled receptor (GPCR)

GNAO1 proteins are known to interact with GPCRs from the GABA, adrenergic, dopamine, and muscarinic group, and others [19], underscoring the diversity and extent of systemic damage incurred with GNAO1 mutations. When activated by their respective agonist, GPCRs act as GEFs for GNAO1.

### GAP: RGS protein

Multiple RGS proteins accelerate the GTPase activity of GNAO1, including RGS1, 2, 4, 6, 7, 9, and 16 [25, 26, 27]. According to The Allen Brain Atlas [12], the most robust coexpression is discernible between GNAO1 and RGS20, 8, 9, 14, and 2 (Table S1), with RGS20, 8, 9, and 2 supported by the analogous data for mouse (Table S2). Out of these, RGS8 has been tentatively linked to seizures in human, while RGS9 knockout mice show dopamine-related motor and appetitive reward deficits [28, 29, 30]. This interaction is possibly competing with the GNAO1-effector binding [11].

### Effectors of G*α* and G*βγ*

The prototypical downstream effectors of GNAO1 signaling are adenylate cyclase for G*α*, and ion channels, phospholipase C-*β*, or phosphoinositide-3-kinase for G*βγ* [19]. GNAO1 functions as an inhibitor of adenylate cyclase (ADCY) *in vitro* [31]. Of all ADCY genes, ADCY5 most convincingly coexpresses with GNAO1 in striatum (Table S3). This interaction is further supported by the analogous co-expression in mouse brain [32], Table S4, and by the recently established connection between mutations in ADCY5 and movement disorders [33, 34].

In parallel, GNAO1’s partner G*βγ* regulates calcium, potassium and sodium ion channels [35, 11]. The potassium channels M and GIRK, in particular, have been implicated, respectively, in the early onset epileptic encephalopathy (EOEE; thought the component peptide KCNQ2) [36] and general epileptic phenotypes in human and in animal models (through GIRK2) [37]. GIRK, in addition, has been reported to be directly regulated by G*βγ* in cardiac myocytes, though the subunit through which G*βγ* regulates GIRK is the heart specific GIRK4, homologous to the neuronal counterparts GIRK2 and GIRK3 [38].

Even with the mediators of signaling downstream of GNAO1 only partly understood, we argue, a predictive power can be gained by considering the GNAO1 itself, and its role as a signal relay. With that aim, we proceed by inspecting the protein location of the disease-causing mutations, and the aspects of the signal transduction they affect.

### Functional regions of GNAO1 and the location of disease mutations

GNAO1 processes almost all of the (inter)actions involved in the GPCR signaling cycle: its central functional part is the catalytic pocket that is normally occupied by either GDP or GTP at any time, and it also interfaces G*β*, GPCR, RGS, and the effector(s). To these functional regions we should add the folding core - buried residues responsible for the proper folding of the protein. GNAO1 must handle all of these through its relatively small structure - even more restrictively, through its catalytic domain (which encompasses but is not restricted to the catalytic pocket, Movie S1). This has three strong implications: the functional interfaces overlap, all residues in the catalytic domain belong to at least one functional group, and all residues have very little freedom to vary. While the latter is the consequence of the heightened evolutionary pressure on residues engaging in multiple interactions with molecular partners, the first two observations follow from the geometry of the problem (Movie S1). In particular, the same interface is responsible for the interaction with G*β* and all of the known effectors of G*α* proteins. This further implies that a single mutation can have impact on multiple signaling pathways, and on multiple points in a given pathway.

### Response to therapy, from the drug target perspective

Our ultimate goal is to be able to understand which therapy, and why, might be applicable for each genotype. As the first step, then, we systematize the therapy response recorded and reported so far (Dataset S1), according to the location of the resulting lesion on GNAO1 protein.

For each drug we extract the activity values toward its main protein targets, which are typically transmembrane receptors (not GNAO1 itself). Drugs commonly target several members of a protein family, and we do not know for certain which members elicit the effect. To make the information more compact and tractable, we group all members of one target family into a single representative target, additionally labeled by the regulation direction (up- or down-regulation; Fig. 1).

**Figure 1:**
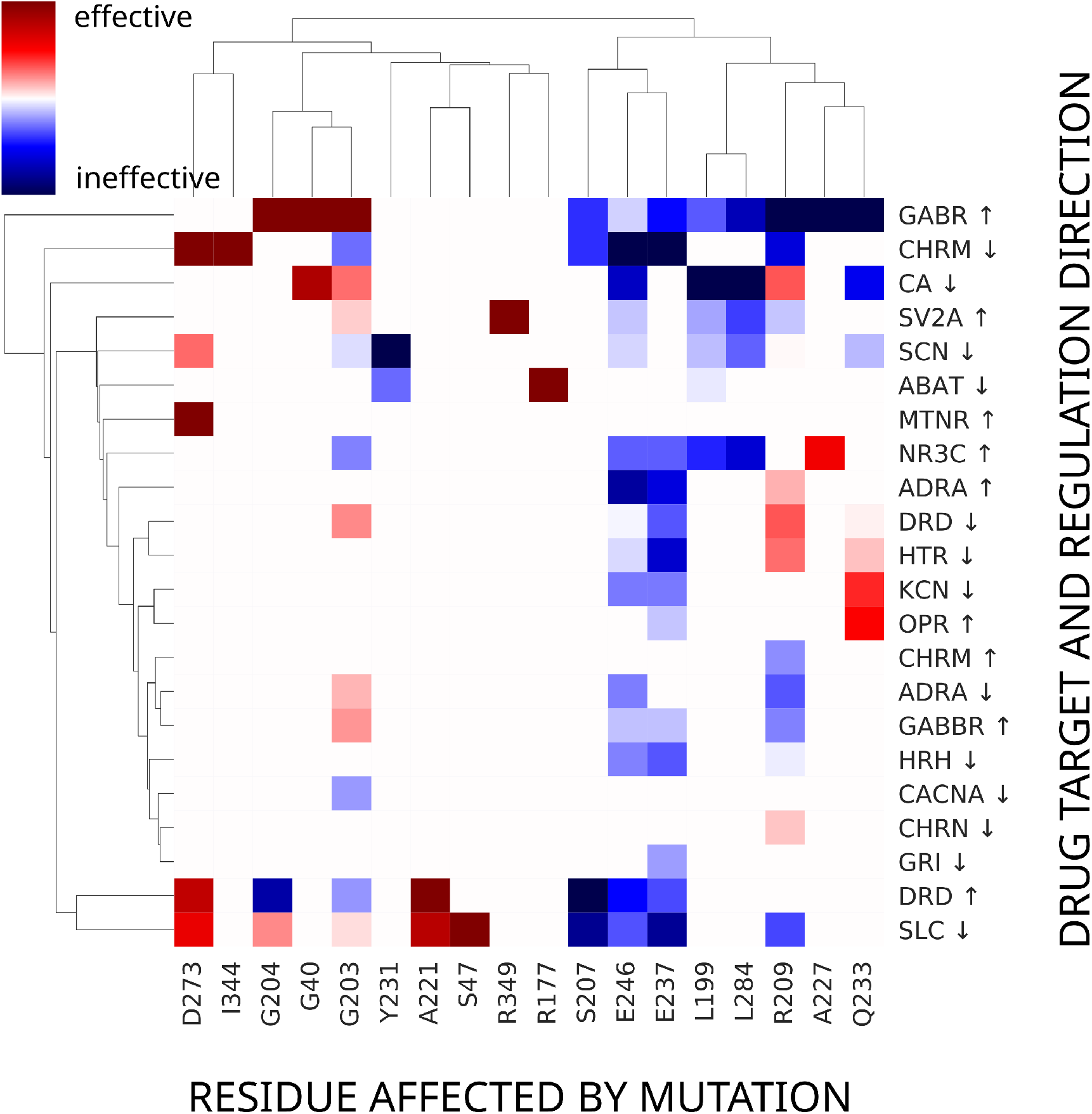
The response profiles for the protein positions of GNAO1 disease variants with reported therapy. Each reported drug is characterized in terms of its main target(s). A drug is termed “effective” if it elicits a response. For most of the patients this does not mean resolution of their symptoms. Drug target family names are standard Human Genome Nomenclature Committee names. See Legend in Dataset S1 for the full list. Most drugs target several members of a gene family, and/or multimeric receptors transcribed from several genes. Particularly discriminating are responses to modulation of GABA, dopmamine and muscarinic actylcholine receptors, as well as synaptic vesicle glycoprotein 2A, and transporters from solute carrier group. Clustering of drug response profiles shows variants falling into two categories: drug responsive and drug unresponsive. The latter category overlaps mostly but not completely with mutations at the interface with the effector (see Fig. 2). Heatmap and clustering by Seaborn [39].

A representative target is assigned a high positive score if it has been targeted with strong affinity, if its effect was documented in multiple patients, and if it has repeatedly been noted as effective in reducing the GNAO1-related symptoms. In the opposite case, when the attempted regulation of the target has repeatedly been documented as ineffective, we assign a high negative score. For details see Supplementary Information (SI) “Building the target-response profiles.” In Fig. 1 the score is represented by the color gradient, blue for negative, red for positive, and the results shown as a heatmap.

The most striking feature in Fig. 1 is the clustering of mutation positions into responsive and non-responsive, in particular to GABA and dopamine-receptor manipulation. In the following we will attempt to rationalize that distinction using the existing knowledge about GNAO1 function.

### Structural map of disease variants and responses to therapy

Next, we collate the information about the therapy response with the structural location of the disease mutations (Dataset S1). As we noted previously [10], epilepsy-related (E) phenotypes usually stem from lesions near the catalytic pocket Fig. 2A, though there is an additional cluster of positions resulting in E phenotypes which cannot be interpreted based on our current understanding of G*α* proteins, Movie S1. Movement disorder (MD) only phenotypes are quite strikingly clustered at the effector interface, Fig. 2B.

**Figure 2:**
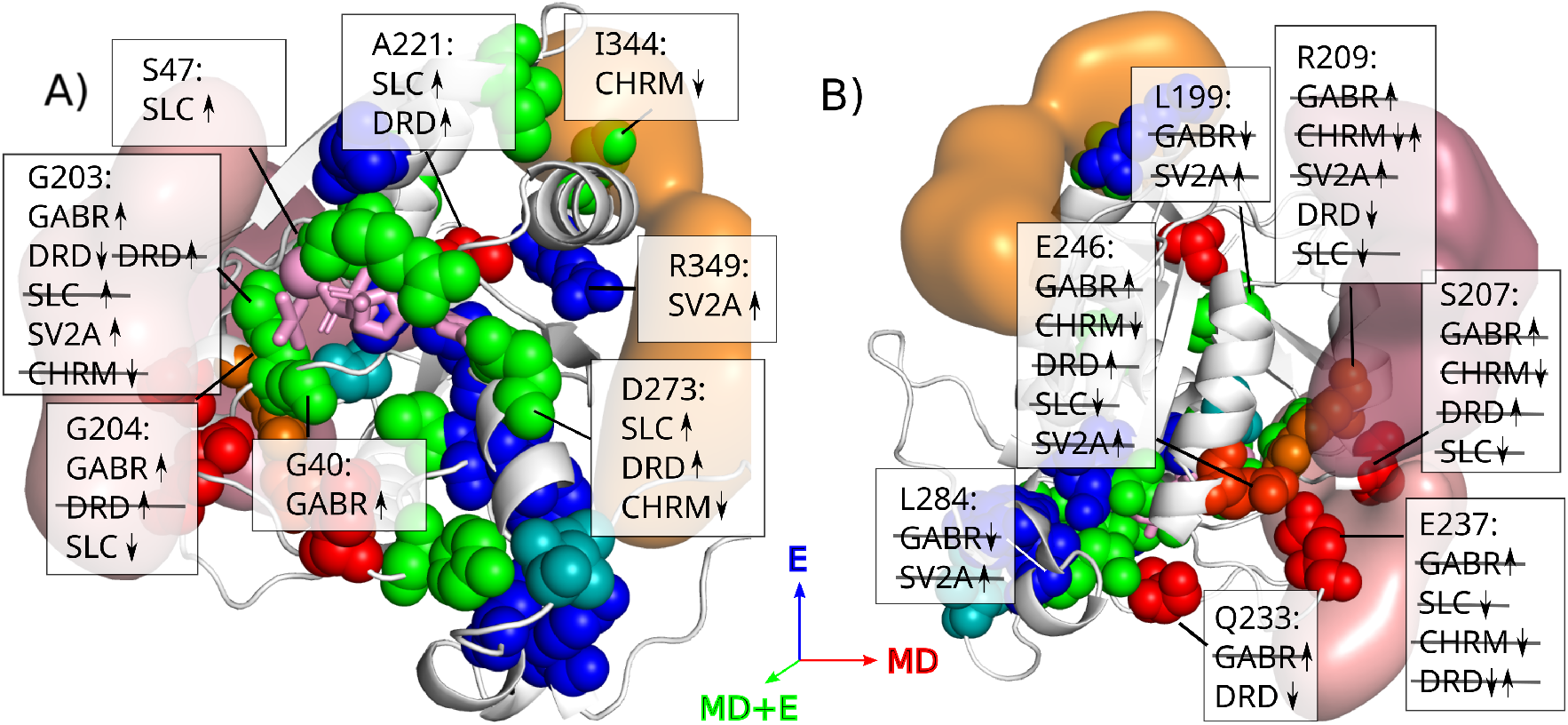
Structural map of disease-related variants in GNAO1, and the main features of the effective therapy reported for their treatment. Orange, surface representation: GPCR. Deep red surface representation: effector. Cartoon representation, white: GNAO1 protein backbone. Sphere representation: position of missense variants (Dataset S1, Legend). Red: movement disorder (MD), blue: epilepsy (E), green: both phenotypes. Cases when different phenotype combinations were reported in different patients are indicated by the combination of the basic RGB colors, Fig. S9. Gene family names same as in Fig. 1. Arrow up(down): up(down)-regulation or increased (reduced) availability. Strikethrough: the therapy shown to be ineffective. This is a low resolution map, in the sense that it ignores subdivision in response possibly introduced by different amino acid replacements at the same mutation position. **A)** View toward the catalytic pocket. Pink: Mg ion; magenta sticks: nucleotide. **B)** View toward the back side of the catalytic pocket, the putative region of interaction with the effector.

### Rationalizing the phenotype through modeling the signal transmitted by GNAO1 mutants

So why do E or E+MD generating mutations cluster around the catalytic pocket and respond to the reported therapies while effector interface, MD-only mutations do not? A qualitative answer is that the mutations in the catalytic pocket, counterintuitively, make the system hyperexcitable. This affects both the G*βγ* and the G*α* branches of the signal. Treatment for G*βγ* entails reducing general neuronal excitability, and, for G*α*, enhancing GABA receptors. The reason for the latter is that because GNAO1 *inhibits* its effector, the hyperactive GNAO1 means hyper-downregulated GABA signaling. Adding to the complexity of the pathway, GABA-ergic neurons expressing GNAO1 may themselves have an inhibitory role [13].

At the other end of the problem, and the protein, none of the attempted MD treatments reported so far address the issue of broken effector (down-) regulation, as is the case when a variant degrades the GNAO1-effector interaction. In the case of ADCY5, if that is indeed the preferred effector of GNAO1, that would entail blocking or downregulation of adenylyl cyclase [40], for which purpose no drugs have been approved so far [41].

We can however develop a better intuition about the systemic effects of GNAO1 mutations by modeling the interrelated reactions within its signaling cycle.

### Modeling the GPCR-related system of biomolecular reactions

We model the GPCR signaling system as a set of reaction rate equations [42], in which the relative concentrations of molecular species (complexes and individual molecules) change in a mutually interdependent way - products of one reaction being the input into another - and all details of the interactions are hidden in the rate constants (Fig. S5). The concentrations [43, 44, 45, 46, 47, 48, 49] and rate constants [50, 51, 52, 53, 54] are chosen to follow as closely as possible the known experimental values (SI, Table S3 and “Modeling agonist dose response”). It is important to note that at this point the model is an explanatory device, rather than a predictive tool, see Discussion below.

### Signal transduction in the wild-type GPCR system

Let us first consider, assisted by the simulation, how the system behaves in its wild-type form. Before the arrival of the agonist signal, most of the G*α* is inactivated by the GDP in its catalytic pocket, and tied in G trimer with G*βγ*. For all practical purposes, the exchange of GDP for GTP never happens spontaneously, and the system is thus paused in the equilibrium state (Fig. 3A).

**Figure 3:**
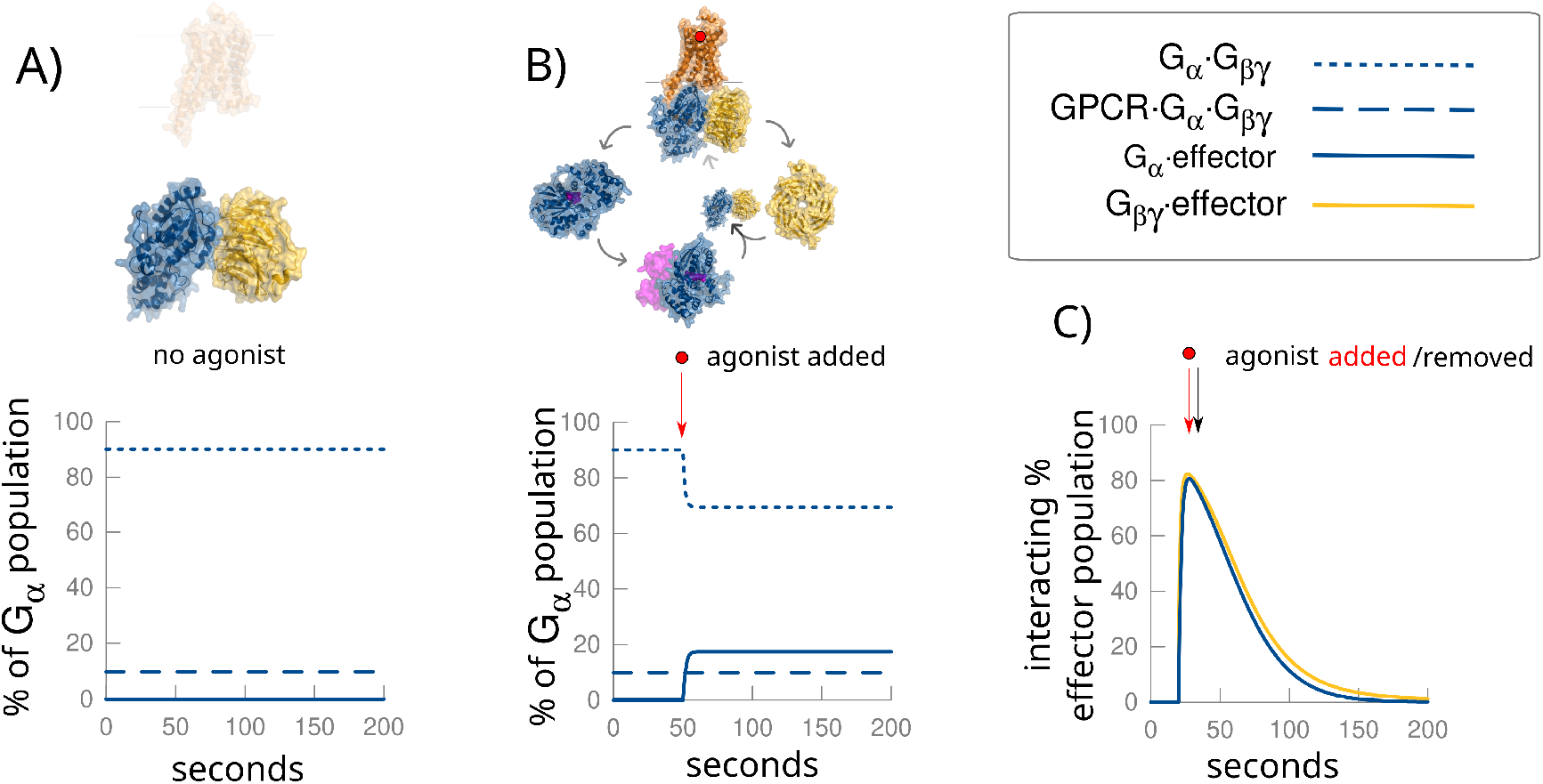
GPCR signal transiently takes the system out of its equilibrium state. **A)** Without agonist present, all G units are bound in trimers. **B)** With agonist, majority the units cycle between trimer, activation by GPCR, interaction with effectors and deactivation with the help of RGS. **C)** Signal: agonist appears, it is removed, and the system returns to its original state. Blue lines: G*α*. Yellow lines: G*βγ*. The interacting partner is indicated by dot notation.

The signal consists in taking the system out of this equilibrium, and letting it relax back (SI, Movie S1). The arrival of the signal, from the perspective of the G-trimer, consists of an active guanine-nucleotide exchange factor (GEF) appearing in the system, with the main effect of enabling the replacement of GDP with GTP. In the GPCR-signaling system, the GEF is the GPCR itself. The ‘appearing’ - in the functional sense - of the GPCR is the allosteric response to extracellular agonist binding. The ensuing GDP for GTP exchange is accompanied by the disassembly of the trimer and its release from the receptor. From this point on, the signal branches in two directions. In the context of this work, G*βγ* is traditionally related to epileptic phenotype (if deregulated), through its interaction with potassium channels, and G*α* to regulating cAMP signaling through ADCY, implicated in movement disorders. The cycle of interactions (not to be confused with the signal) is closed with G*α*, assisted by GAP, catalyzing the GTP hydrolysis, followed by the reassembly of the G-trimer.

Each of these interactions is an ongoing process that repeats as long as the participating entities are available. Thus, the overall effect of an active GEF is the repartitioning of the available G*α* and G*βγ* into an active population, consisting of G*α* and G*βγ* bound to their respective effectors, and the populations of free and GPCR bound G-trimers (Fig. 3B). The relative size of each molecular population depends on the ratio of interaction rates and relative abundance of molecular species (SI, Fig. S5).

Once the signal is terminated (by removal of the GPCR agonist) the system begins its return to the original, equilibrium state. Thus a brief salvo of agonist molecules results in an intracellular signal extended in time, of the characteristic shape [55], shown in Fig. 3C. The GPCR-G signal decays on the timescale of seconds to tens of seconds [56, 57], depending on the mechanism through which the agonist appears and is removed from the system [14]. In our model we use an acetylcholine-like mechanism, in which the agonist molecule is removed by the action of acetylcholinesterase (AChE). AChE cleaves the agonist and thus removes it from the system. This should be understood as a stand-in for any agonist removal mechanism that may be used by a GPCR system (such as dopamine reuptake). The particular choice may modify the timescale but not the signaling cycle itself.

As a measure of the strength of the signal traveling along each of the two branches we use the fraction, or percentage, of G*α* and G*βγ* effectors engaged in interaction with G*α* and G*βγ*, respectively, Fig. 3C. These interactions are assumed to result, for example, in the regulation of ion current by G*βγ* and ADCY inhibition by G*α*, which are not explicitly modeled.

### Signal modification by changing reaction rates

The height and the shape of the signal depend on the catalytic rate of GNAO1, as well as its affinity for the other molecules in the core signaling cycle. These properties – in our model reflected in the reaction rates (Fig. S5) – can be modified by mutations in the peptide, and depend on the physical properties of the replacement (for abundance, see below). In principle, each mutation will have some effect on each of the interactions in the system. However, the relative size of the impact on various interaction interfaces is expected to depend on the distance/proximity to each of them. When designing a decision platform for precision medicine, the change in reaction rates should be either evaluated in molecular simulation, or preferably determined experimentally. We choose to limit our assessment of mutations to those that *weaken* an interaction, and observe signal propagation through the system given the reduced interaction rates. In all of the clinical cases reported so far, the mutations were *de novo* and dominant; therefore, in our simulations only half of the G*α* population is mutated. The details for the mutations discussed in the following sections can be found in Supplementary Dataset S1.

#### Null mutants

First, let us note that as long as GNAO1 outnumbers its GPCR and effector partners, null mutants, the only effect of which is to halve the GNAO1 population, have no effect on the GCPR signal, Fig. S6. The null mutants in this context might be either early stop codons (nonsense and frameshift mutations), or mutations causing protein misfolding. Consistently, no early stop codon mutations were reported in GNAO1 patients, while gnomAD v.2.1.1 [58] reports one healthy person heterozygotic in L269 to nonsense mutation, a position sufficiently early in the sequence for the elimination of the transcript by nonsense mediated decay.

#### Mutations affecting GTP to GDP catalysis

The most commonly seen dysfunction of GNAO1 mutants can be understood in terms of their reduced catalytic capability. This may be the result of degraded interaction with the activating protein (RGS; Fig. 4A), or the alteration of the catalytic pocket (directly or remotely; Fig. 4B).

**Figure 4:**
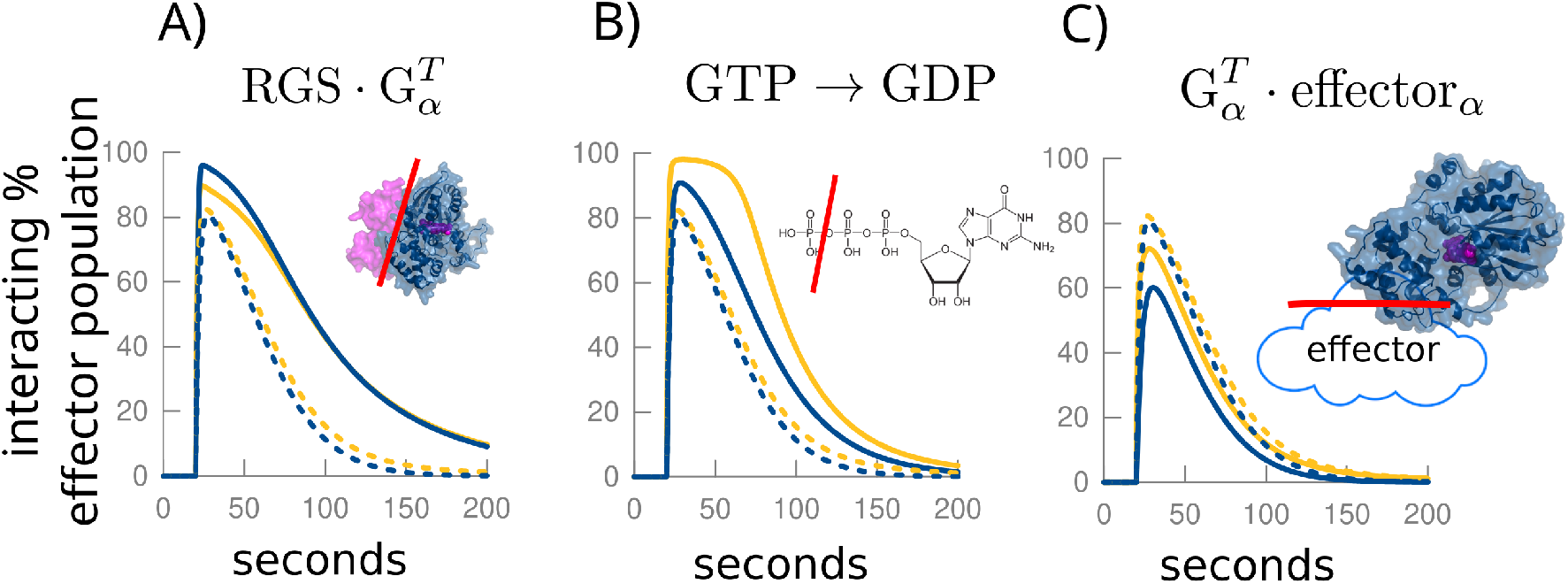
Signal modification. By mutation that **A)** changes the interaction with RGS, **B)** alters the efficiency of GTP to GDP catalysis, and **C)** changes the interaction with the effector. Blue lines: G*α* signal. Yellow lines: G*βγ* signal. Dashed lines: wild type signal. Solid lines: modified signal. The interacting partner is indicated by dot notation. In all cases the mutation is assumed to reduce the forward rate of interaction. Note that the only way for a mutation to downregulate the interaction with the effector is to exclusively modify the interaction with its interface, which is much less likely than having a mutation with multiple effects.

Many positions mutated in disease are in the close proximity to the catalytic pocket (G45, S47, R177, N270, D273) all closer than 4*Å* to the substrate; Dataset S1, and given the precision required for the process, it is reasonable to assume that as a result the catalysis is degraded, rather than being enhanced by chance. Additionally, keeping in mind that the catalytic domain is relatively small, it is conceivable that even a mutation not directly within the catalytic pocket can distort its geometry, and exert a similar effect (I56 and D174, tentatively). A slowed GTP catalysis means that G*α* is stuck with the GTP bound, and as long as GTP is present, G*α* is primed to interact with the effector, resulting in extended signal. The increased G*α* activity means increased *downregulation* of the effector, which seems to result in MD phenotypes that do not include chorea or ballism. Rather, the phenotypes include severe hypotonia, myoclonic seizures, spasticity, sometimes dystonia.

G*βγ* branch follows G*α* closely in this case, Fig. 4A. We would expect, thus, that both sets of symptoms react well to a single therapy consisting of reduction of the availability of the agonist, damping down the signal. This is admittedly difficult to discern from the available case descriptions in the literature. The patients *were* reported (Dataset S1) to respond to reduction of monoamine neurotransmitter availability (tetrabenazine) and GABA replacement (vigabatrin). In the cases of extreme dysfunction of the catalytic pocket, no therapy may be effective (“Exotic effects” below; Dataset S1).

#### Mutations affecting the interaction with the G*α* effector

The number of ways in which the G*α* signal can be modified exclusively is small. Either direct degradation of the effector interface, or, sometimes, a chance superposition (see below) can generate the effect. Our best candidate for the mutation positions that result in the degraded effector interface is S207. If the interface with the effector is degraded, we expect a signal of reduced amplitude, though not necessarily of significantly shortened duration Fig. 4C. The problem, from the therapeutic perspective, is that the result is an intensified effector signal, presumably cAMP, for which no approved downregulator exists. The reported phenotypes include severe choreoathetosis, dystonia, and ballism, difficult to treat even in the intensive care, Dataset S1 and the references therein. Mostly poor or no response to dopamine and GABA manipulation has been reported in these cases, Fig. 1.

#### Double impact mutations

Because the catalytic domain of GNAO1 is relatively small, it is feasible for a mutation to have impact on multiple aspects of its function. In particular, some of the experimentally available data [43] can be modelled by assuming that some mutations have double impact, simultaneously affecting the catalytic pocket and the effector binding interface (SI, “Modeling agonist dose response”). Note that this mechanism is different than mutants exerting impact on both the G*α* and G*βγ* signaling branches. Both branches are always affected, with very little exception. Independently of that effect, the shape of the distorted signal depends on which G*α* interactions are degraded. For example, while a mutation that slows down the catalysis intensifies and prolongs both branches of the signal, Fig. 4A, a double impact mutation may suppress the G*α* branch, while enhancing the G*βγ*, Fig. 5A. A straightforward example of the latter is provided by mutations in G203 and G204, less than 6*Å* from GDP and 8Å from effector, that result in both EOEE, and severe choreoathetotic symptoms, but without the severe hypotonia that we see in the context of the (exclusive) degradation of catalysis efficiency.

**Figure 5:**
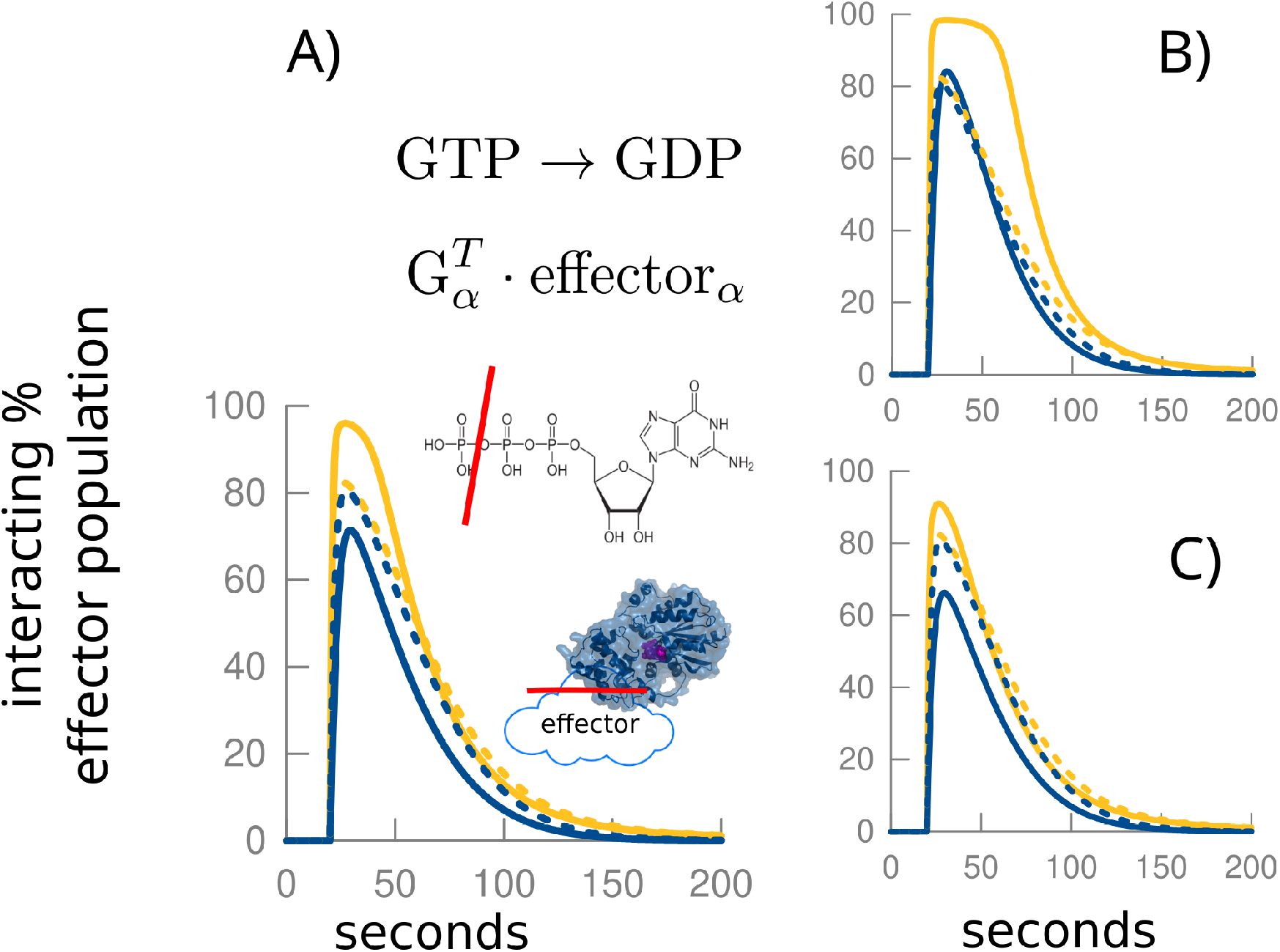
Compensating effect of double-impact mutations. A simulated effect of a mutation having a simultaneous impact on the catalysis and effector binding. Blue lines: G*α* signal. Yellow lines: G*βγ* signal. Dashed lines: wild type signal. Solid lines: modified signal. **A)** In this hypothetical case the G*α* signal (blue) is suppressed, and G*βγ* (yellow) is increased. However, **B)** a small increase in the catalysis rate can make G*α* branch behave almost like the wild type (note the overlap of the blue solid line with the dashed lines), while **C)** a small decrease can produce analogous effect in the G*βγ* branch (the overlap of the yellow solid line with the dashed lines).

A particular consequence of double impact mutations is that they may result in superposition of two effects moving the signal in two opposite directions (enhancement vs. suppression). In extreme cases and in the right environment (relative concentration of agonist, GPCR and the effector) this may lead to their mutual compensation, with the resulting signal for one branch close to its wildtype shape, Fig.5B.

A prototypical example is provided by G40, a residue buried deep inside the structure, some 6*Å* away from the substrate and 9*Å* from the effector. Several cases have been reported, all with bulky amino acid replacements (G40W, G40R, G40E). The behavior of the G40R mutant under conditions measured by Feng *et al.* [43] can be modelled well under the assumption that it produces sizable effects on both interfaces, see SI, “Modeling agonist dose response”. The distance from G40 to the catalytic pocket and the effector is slightly larger but comparable to the cases of G203R and G204R, where hyperkinetic movements are prominent, Dataset 1, yet the movement disorders recorded for G40 were less robust (either non-existing, limb twitching, or a single limb dystonia [10]). We suggest that this is the result of the effector interface being degraded just to the extent that can be compensated by the concomitant slowing down of the catalytic process, which extends the time in which G*α* is available for the interaction.

Another extreme example of this effect is provided by the mutations in E246 and R209, E246K and R209H in particular. In the experiments by Feng *et al.* they are indistinguishable from the wild-type. Even more strikingly, gnomAD v2. database records four heterozygote individuals with E246K mutation that are apparently healthy [58]. This time, in the model at least, it is possible that the ratio of rate constants is moved in the other direction, Fig.5C, masking the impact on the catalytic pocket brought about by disruption of the salt bridge between E246 and R209, exposing only the severe movement disorder phenotype.

#### Mutations affecting the interaction with the GPCR

The disease mutations involving the G*α* C-terminal helix through which it interacts with GPCR are rare (I344del, R349-G352delinsQGCA), and the phenotype relatively mild (seizures limited to early childhood, staring spells; mild spasticity, oromotor apraxia). A single mutated allele with decreased affinity for GPCR should not be able to produce a noticeable effect Fig. 6A if the ratio of GPCRs to G*α* is small (see SI, “Parametrizing the GPCR signaling system”). Even when that is not the case, when the ratio is 1:1, the effect is small, Fig. S8, and the reported cases responded to trihexyphenidyl, usually thought of as anticholinergic. We ascribe that effect to the often quoted [59, 60] but seldom investigated [61, 62, 63] possibility that the role of trihexyphenidyl is mediated by enhancing the availability of dopamine. According to Shin *et al.* [63], activation of muscarinic M2 and M4 acetylcholine receptors (CHRM2 and CHRM4) in the nucleus accumbens depresses, whereas activation of CHRM5 potentiates dopamine transmission, Fig. S3. Trihexyphenidyl, on the other hand, is a CHRM *antagonist* with the affinity of 1-3nM for CHRM4, 7-12nM for CHRM2, and and 9-16nM for CHRM5 [64]. In our model, enhancing the availability of dopamine would enable the system to overcome the weak baseline signal.

**Figure 6:**
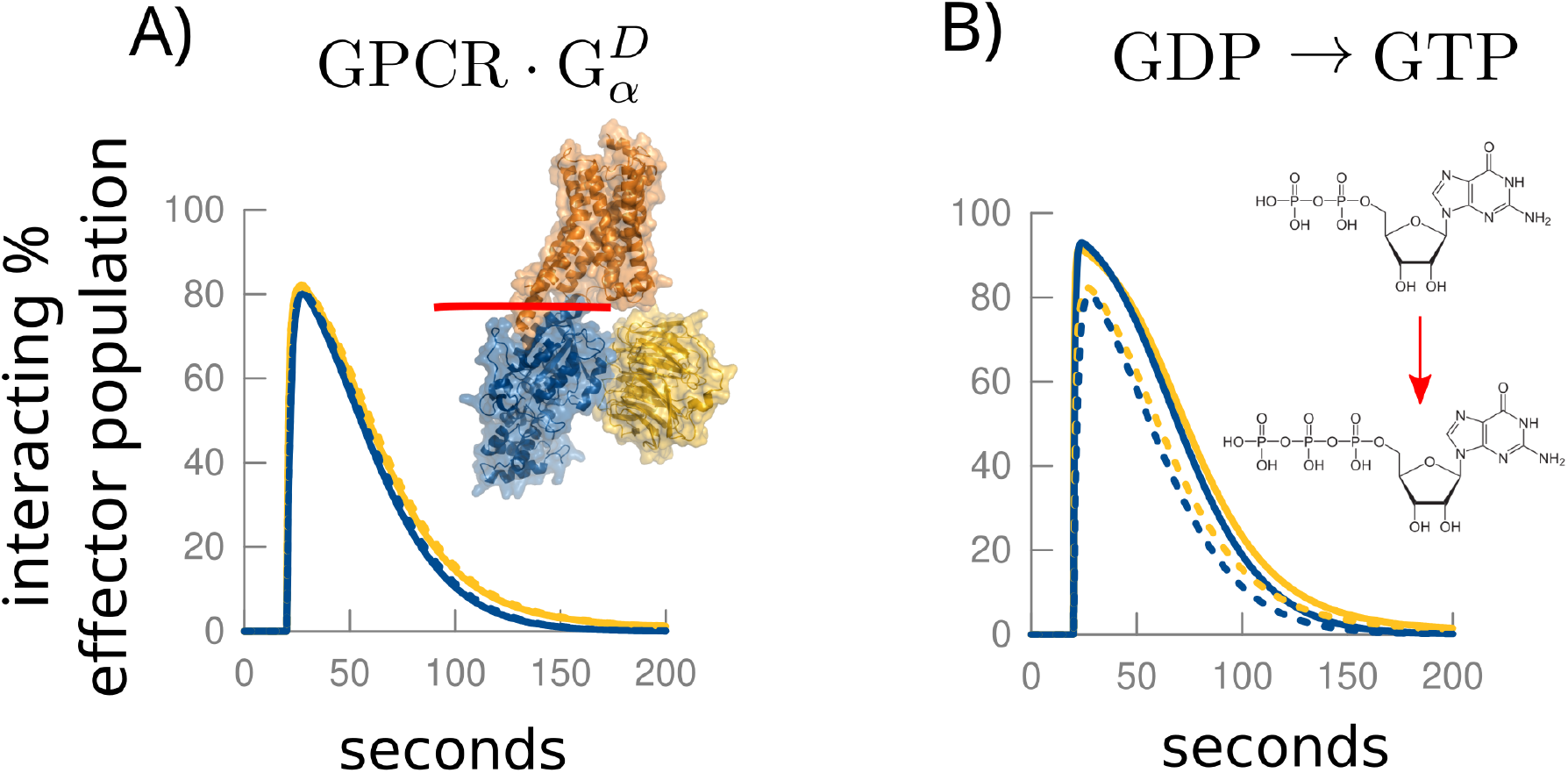
Impact of mutations at the GPCR interface. **A)** Effect of weakening the interaction between G*α* and the GPCR is nonexistent. In the system in which two alleles compete for a much smaller number of GPCRs, the wild-type allele outcompetes the mutant. **B)** xIf, however, the mutation enhances the GDP for GTP exchange, the effect is a notable increase in the signal amplitude. Dashed: wild type.

It is possible that the effect of the R349-G352delinsQGCA is exerted through a more subtle change in the function of the protein (see SI, “Forme fruste of GNAO1 phenotype”) leading to the enhanced GDP for GTP exchange. This, in the model, leads to increase in the signal peak intensity Fig. 6B. The patient responded well to levetiracetam, believed to reduce the availability of the agonist.

#### Epilepsy cluster

When discussing the effector-only impact mutations, an important note should be made of the cluster of positions away from all known G*α* interacting interfaces, Movie S1, mostly causing epilepsy-only phenotypes (peptide positions 231, 275, 279, 284, 291, and possibly 228), Movie S1. These spatially distinct mutations suggest a function that is specific to GNAO1, which means that it is probably *not* related to folding and structure stability (though it should be noted here that Y231C, F275S, and I279N did not express well when transfected into human embryonic kidney cells in the experiments of Feng *et al.* [43]). Another possibility is that the effect of these mutations is related to an interaction interface specific to GNAO1. Indeed, that region is conserved in all vertebrate GNAO1s. Taken together, these observations suggest the possibility of so-far unrecognized GNAO1 effector(s), possibly in different cell types, leading to epilepsy phenotypes not mediated by G*βγ*.

#### Exotic effects: empty catalytic pocket

Some mutations may lead to special behavior, completely abolishing some or multiple interactions in the GPCR signaling cycle. The case in question here is G42R mutation, presenting a unique combination of severe hypotonia and choreoathetosis, the first one, in the majority of the cases we have collected, is associated with extended G*α* signal, and the latter with the lack thereof. In Fig. S7 we show the behavior of the “empty pocket” mutant - a Go*α* mutant that does not bind its substrate-described in the series of works by Yu *et al.* [52, 53, 54]. It has two peculiar properties of blocking its native GPCR, even in the absence of the agonist, while not binding its G*βγ*. Depending on the ratio of concentrations of G*α*, GPCR and the effector in the system, these properties can lead to permanently elevated baseline signal for the G*βγ* and almost completely suppressed G*α*, the latter corresponding well with the severe choreoathetosis. G42R is a good candidate for this model because the bulky substitution should completely displace the last phosphate group in the nucleoside making both the binding and the catalysis very unlikely. To move this argument forward, we need to learn how the hypotonia arises in this context in the first place.

## Discussion

We have argued that the continuum of GNAO1 phenotypes is the consequence of the continuum of changes that a mutation can produce in this protein. Initially, one might expect that a mutation having an impact on a single aspect of GNAO1 function should manifest as a single well-defined phenotype. There are at least two reasons why that is not always the case. One, because the GPCR signaling cycle branches and closes back onto itself, and disruption in any interaction changes the availability of the reactants for the remaining interactions, it is almost impossible to change one branch (G*α*, MD-related, or G*βγ*, E-related) without affecting the other; and two, because the small and compact geometry of the protein practically prevents mutations from having an isolated effect on a single function.

Consider the mutations that exclusively affect the catalytic pocket. They are the closest to having a clear-cut, single-function impact on GNAO1 protein. Indeed, they universally result in early-onset epileptic encephalopathy [10]. However, the milder, non-choreoathetotic MD symptoms are present concomitantly, because on the system level, the branched signal that does not withdraw properly, Fig. 4B, creates prolonged impact on both G*α* and G*βγ* effectors. Accordingly, the patients with these variants seem to react to lowering neuronal excitability or neurotransmitter availability. Increasing GABA-sensitivity of the downstream neurons also makes sense in the context of striatum, since we are reasonably sure that GNAO1 there functions as a downregulator of ADCY5 in GABAergic neurons.

In contrast to the catalytic pocket mutations, the majority of GNAO1 mutations have an impact on multiple functional sites. If a distant mutation, i.e. a mutation not in the immediate vicinity of any of the functional sites, weakens the reaction with the effector, the change can be compensated for by increasing the availabilty of the neurotransmitter. We can begin to see, though, the conflicting requirement that it poses - the distant mutation is likely to have a simultaneous impact on the catalytic pocket, making the G*βγ* branch hyperactive and in the need of neurotransmitter *reduction*.

When the mutation completely abolishes the interaction with the effector, as might be the case with some mutations exactly at the interface, the resulting MD phenotype is refractory to all reported therapies. We suspect the reason is that the available drugs do not address the underlying problem - the broken downregulation of the effector. As these mutations disconnect GNAO1 from the effector, they cannot be treated by enhancing the GPCR cycle. Therapies must therefore downregulate the effector directly. Yet, even for the prototypical effector, ADCY5, the needed downregulator is not available, as of this writing (though agents exist that might achieve this purpose [41]).

How do we, then, select therapies for novel GNAO1 variants? Since the phenotypes of various molecular lesions overlap greatly, in particular in our limited ability to describe them, it is unlikely that we will ever be able to discern the impact on the molecular level just by observing the phenotype. Even the mapping of a genetic variant onto protein structure proves to be insufficient, because of the indirect impact it may have through distorting the structure. A dedicated experimental pipeline for quantitative characterization of GNAO1 mutants, evaluating their level of expression and reaction rates with all interacting partners, might be of help to the growing number of patients diagnosed with GNAO1-related disease. Coupled with better understanding of relative abundance of molecular members of the GPCR signaling cycle in the relevant types of neurons, a computational model, such as the one used in our discussion here, would become a predictive tool useful in choosing the first line of drug therapy.

But even in a hypothetical case where we can perfectly quantify the impact of a mutation on each of GNAO1’s functions, we are still left with the problem of specifically targeting the system that as its input and output has such generic entities as dopamine and GABA. Looking ahead, it might prove cost efficient in the near future to move to a gene therapeutic approach.

GNAO1-related diseases are caused by autosomal dominant variants. Our model suggests why: ultimately, the mutants exert their effect by competing nonproductively for the resources in the the signaling cycle. It may seem counterintuitive, then, that in the healthy population there are many variants causing nonsynonymous mutations in the highly conserved catalytic domain of GNAO1 [58], Movie S1. An important lesson can be drawn from null variants, the variants that completely remove a GNAO1 allele from the pool of expressed proteins. While there are several such variants reported in the healthy population, none are reported in the context of the GNAO1 disorders. A variant that causes misfolding on the protein level may have a comparable systemic effect, or rather, lack thereof. Removed from the competition for resources, such alleles are neutralized from producing pathological phenotype. This points to a straightforward path to gene therapy in GNAO1-related diseases: silencing of the faulty copy of the gene. Several anti-sense oligonucleotides are already in clinical trials for neurological diseases [65], in particular for the treatment by gene expression downregulation [66, 67], and we expect CRISPR-based technologies to follow shortly [68]. Understanding the impact of a variant on the expressed protein will then become crucial in choosing a personalized approach to GNAO1 patients.

## Materials and methods

### GNAO1 splice

All mutations discussed in this work refer to the GNAO1 splice most commonly expressed in the brain, NCBI accession number NM 020988.2.

### Molecular structures - modeling and visualization

The model for active GNAO1 protein structure was built on the template from PDB [69] identifier 3c7k [27], using Swissmodel [70]. The same PDB entry was used to model the interaction between G*α* and and RGS domain. The interaction with GPCR modeled after PDB entry 3sn6 [71], and interaction with acetylcholinesterase after 6r3q [72]. Mapping between various complex structures was done using deconStruct [73]. The illustrations were generated using PyMol [74].

### Modeling GPCR-G signal

The set of equations describing the core GPCR signaling cycle, Fig. S4, was solved using BioNetGen [75]. The input for BioNetGen was initially prepared using V-Cell [76]. The parametrization was taken from the published biochemical experiments [52, 53, 51, 50], details: Fig. S5.

## Supporting information

Supplementary text, figures and tables. Includes model parametrization.

## Code availabilty

The full analysis pipeline used in this work can be found in GitHub repository (github.com/dogmaticcentral/gnao1), and the code needed to reproduce figures in the text in the compute capsule on CodeOcean (codeo-cean.com/capsule/8747824).

