## Supplementary text, figures and tables. Includes model parametrization. for "Molecular map of GNAO1-related disease phenotypes and reactions to therapy"

#### This PDF file includes:

Supplementary text
Figs. S1 to S9
Tables S1 to S5
Legend for Movie S1
Legend for Dataset S1
SI References

#### Other supplementary materials for this manuscript include the following:

Movie S1
Dataset S1

### Supporting Information Text

### Building the target-response profiles

To construct the target-response profiles used in Fig. 1 in the main text, we collected the information about the targets of drugs reported as therapy for GNAO1-related symptoms. We collected both the direction of action (up- or down-regulation), and its micromolar activity for each drug-target pair, keeping in mind that drugs typically have multiple targets. The sources were DrugBank (1), BindingDB (2), GuideToPharm (3), PDSP (4), and PubChem (5), as well as manual literature search in a handful of cases. This information is included on Dataset S1. The full collection can be found and downloaded as an SQLite database from the accompanying CodeOcean capsule (codeocean.com/capsule/8747824).

Giving the name $td$ to the combined label of target + drug activity direction, for example $GABR\uparrow$ for upregulated GABA-A receptor (Human Genome Nomenclature Committee symbol GABR), we assign to it a weight

$$w(td) = \begin{cases} -log_{10}(\text{activity}) + 6, & \text{if the activity is } < 10^6 \mu M \\ 0 & \text{otherwise.} \end{cases}$$

This then enables us to assign the profile $W$ to each protein position carrying a disease mutation, for example A221, as

$$W(\text{position}, td) = \frac{1}{N} \sum_{patients} \left[ w^e(td) - w^i(td) \right]$$

The sum runs over all patients reported as carrying the variant that results in the missense mutation at this position in the protein. The $e$ and $i$ superscripts refer to whether the drug was reported to exert a positive effect on the patient (effective; $e$) or not (ineffective; $i$). The norm $N$ is inserted there to make all profiles scale on the range $[-1, 1]$. That is, $N$ is the maximal absolute value for any $w(td)$. In the Fig. 1 in the main text the $[-1, 1]$ scale is replaced with the color representation ranging from blue to red, as indicated by the colorbar. If several drugs were reported to have been used in the same patient, the one with the greatest activity toward the target $td$ was chosen.

To simplify the presentation and compensate for the fact that we are working with a small dataset (from statistical perspective) all target genes were grouped into families and the highest known activity used. In a larger dataset it would be preferable, of course, to look at individual gene targets, and also at replacements on the protein level (A221D, for example) rather than positions themselves.

### Parametrizing the GPCR signaling system

**Stoichiometry of the GPCR signaling system.** Modeling the system requires several quantitative assumptions, or parameters, as the input. In particular, we need a reasonable estimate of the relative abundance ratios of different components in the GPCR signaling system.

The stoichiometry of the GPCR signaling system has been a matter of debate, Ostrom (6), for example, put the ratio of receptor:G protein:effector to 1:100:3, in the system of $\beta$-adrenergic receptor signaling through ADCY via G$\alpha$ of type "s" (GNAO1 is of the type "o"). However, there is increasing evidence that the GPCR signaling machinery does not diffuse freely through the membrane, but that the receptors form homo- and heterodimers (with coexpressing GPCRs from the same family) (7, 8), perhaps further organizing into 2:2 receptor heterotetramers (9), and higher order paracrystalline arrays (10). Furthermore, the results of Nobles (11) show that GPCR dimers can pre-form pentameric associations with G-protein trimers, even in the absence of the agonist. This suggests the 2:1:1 ratio for receptor:G protein:effector. It can be seen, by comparison of Figs. 4 and 6 in the main text, produced using the 1:10:1ratio, and the Fig. S8 here, using the 1:1:1 ratio, the main features of the signal do not depend on that choice. However, the experiments of Feng *et al.*, below, can be explained more naturally using the former.

Since the cell maintains an abundant level of GTP (12), we take that GTP is always and instantly available, and not a rate limiting step in any of the interactions (13).

**GPCR pathway, a biochemically well studied system.** The other big group of input parameters are the forward and reverse rates for the biochemical reactions in the GPCR pathway. Our ability to build a quantitative model of GPCR system owes to many decades of its intense biochemical investigation (13–15). For the full set of parameters used as default (wild-type) system, see Fig. S5.

Of particular interest here are G$\alpha$ mutants that *do not* bind nucleotide (15–17), Fig. S5, lower right. They have two peculiar properties of blocking its native GPCR, while not binding its G$\beta\gamma$. These properties result in a unique distortion of the GPCR signal (Fig. S8, 'empty pocket'). From experiments of Yu and Simon (16), we take that the mutants that do not bind GTP or GDP (such as $G_o\alpha$D273N reported therein) can bind the receptor in the absence of G$\beta\gamma$ and the agonist, but are never (or rarely) released. The kinetic constants for the interaction between the receptor and the empty mutant or its surrogate $G_o\alpha$X (a double mutant GNAO1 regulated by Xanthine nucleotide), were not explicitly reported in this work. However, from the overall similarity to behavior of G-protein heterotrimer that includes GNAO1, we took the assumption that the forward binding rates to GPCR are comparable to wild-type GNAO1. The same lab also reported that $G_o\alpha$X, will not bind G$\beta\gamma$ in the absence of XDP, xanthine diphosphate (15), and we thus also take that the impaired nucleotide binding implies impaired binding of G$\beta\gamma$ in any $G_o\alpha$ mutant.

### Modeling agonist dose response: implications on understanding the impact of disease variants

Important source for the development and parametrization of the model we are using in this work have been the experiments by Feng *et al.* (18). Reproducing the behavior of the mutant GNAO1 system was one of the main constraints in determining the relative abundance of molecular species in the system.

In the setup described by Feng *et al.* in (18), a competing species of $G\alpha_s$ is present in the system. It binds weakly to the same GPCR, and *stimulates* the same adenylyl cyclase, leading to severalfold increase in the cAMP signal when GNAO1 is not present. If we introduce $G\alpha_s$ in our model with suitable parametrization, Table S5, we can reproduce that behavior. Fig. S1 shows the main features of Feng *et al.* Fig. 2 (outlined in the inset) reproduced in the simulation.

Varying the catalytic competence of GNAO1 replicates the behavior from Fig. 2C in Feng *et al*, Fig. S1A here. The candidate positions for the mutations generating that behavior include G45, S47, R177, N270, and D273, see discussion in the main text ("Mutations affecting GTP to GDP catalysis"). G42R mutant, we believe, based on the phenotypic characteristics of the patients carrying that mutation, might actually be a case of an extreme malformation of the catalytic pocket (see "Exotic cases" in the main text).

Reducing the interaction with the interface leads to behavior shown in Fig. S1B, Fig. 2B in Feng, corresponding to mutants R209G, R209C, as well as the variants resulting in the mutated S207.

The most interesting behavior stems from mutants that can disable both the catalytic pocket and the interface with the effector ("Double impact mutations," main text), Fig. S1C, left. This leads to behavior that mimics the non-existent GNAO1 (red curve) in the Feng *et al* setup. However, it should be kept in mind that in the same setup the same effect can be explained in large part by the reduced expression of GNAO1 mutants, Fig. S1C, right. It is reasonable to expect that mutations that affect the folding have both traits: they reduce the folding efficiency, and result in misshapen protein structure that distorts the catalytic pocket, as well as the effector interface.

As we have seen in the panels A and B of the Fig. S1, reducing the rate of catalysis and degrading the effector interface move the response curve in the opposite direction. Thus, as we may expect intuitively, there is a regime of parameters which closely mimics the wildtype behavior in this experiment, Fig. S1D. It is interesting to note that this is precisely the case with the E246K mutant that appears in several healthy people, as reported so far.

As the reduction in the functionality of both sites decreases, we are approaching the no-GNAO1 case. However if these interactions are completely abolished, perhaps counterintuitively, the mutant may overshoot the no-GNAO1 case in its proficiency, Fig. S1E, which can be explained by resource competition - the crucial resource here being RGS - without RGS available the $G\alpha s$ stays activated longer resulting in stronger cAMP current. Indeed, by introducing more RGS and GPCR in the system we can reduce that effect, though the overall behavior then does not correspond to the one described in Feng et al setup.

The code used in generating the Fig. S1, including the parametrization for each case, can be found on CodeOcean (codeocean.com/capsule/8747824).

### Forme fruste of GNAO1 phenotype: an example

In some cases the phenotype itself appears ambiguous. The matter is further complicated by the uncertainty in the assessment of the impact of the disease causing variant. For example, we have encountered and described (19) a case of a curious combination of 10 nucleotide insertion and deletion, leading to two amino acid replacements - R349Q and G352A - in the GPCR interacting helix of $G\alpha$. The effect of the two should be subtle, since the replaced chains are facing away from the GPCR in the interacting configuration. However, in the GTP bound closed conformation, R349 is stashed in a salt bridge formed with E318 (Fig. S2A). This interaction is conserved (Fig. S2C), and may promote the dissociation from GPCR upon GTP binding. Conversely, R349Q has the salt bridge weakened, possibly leading to faster reorganization of the protein upon docking to GPCR, and leading to faster GDP-for-GTP exchange.

But then, again, the GTP/GDP exchange can be increased one hundredfold and yet result in the minuscule increase in the signal. Furthermore, this increase can be rescued by a 10% reduction in agonist concentration (Fig. S2B). Consistently (though possibly unrelatedly), the patient's symptoms are relatively mild, with atypical staring spells and abnormal EEG suggesting seizures, and oromotor apraxia as her only motor disorder (19). She was treated with levetiracetam for one year, and the episodes resolved during the treatment, without re-occurrence after the treatment was stopped. levetiracetam is an agonist of the synaptic vesicle protein 2A, SV2A (20), which is thought to inhibit neurotransmitter release (21).

The fact that the patient reacted well to levetiracetam administration is consistent with the tentative ability of this mutant to overreact to agonist signal.

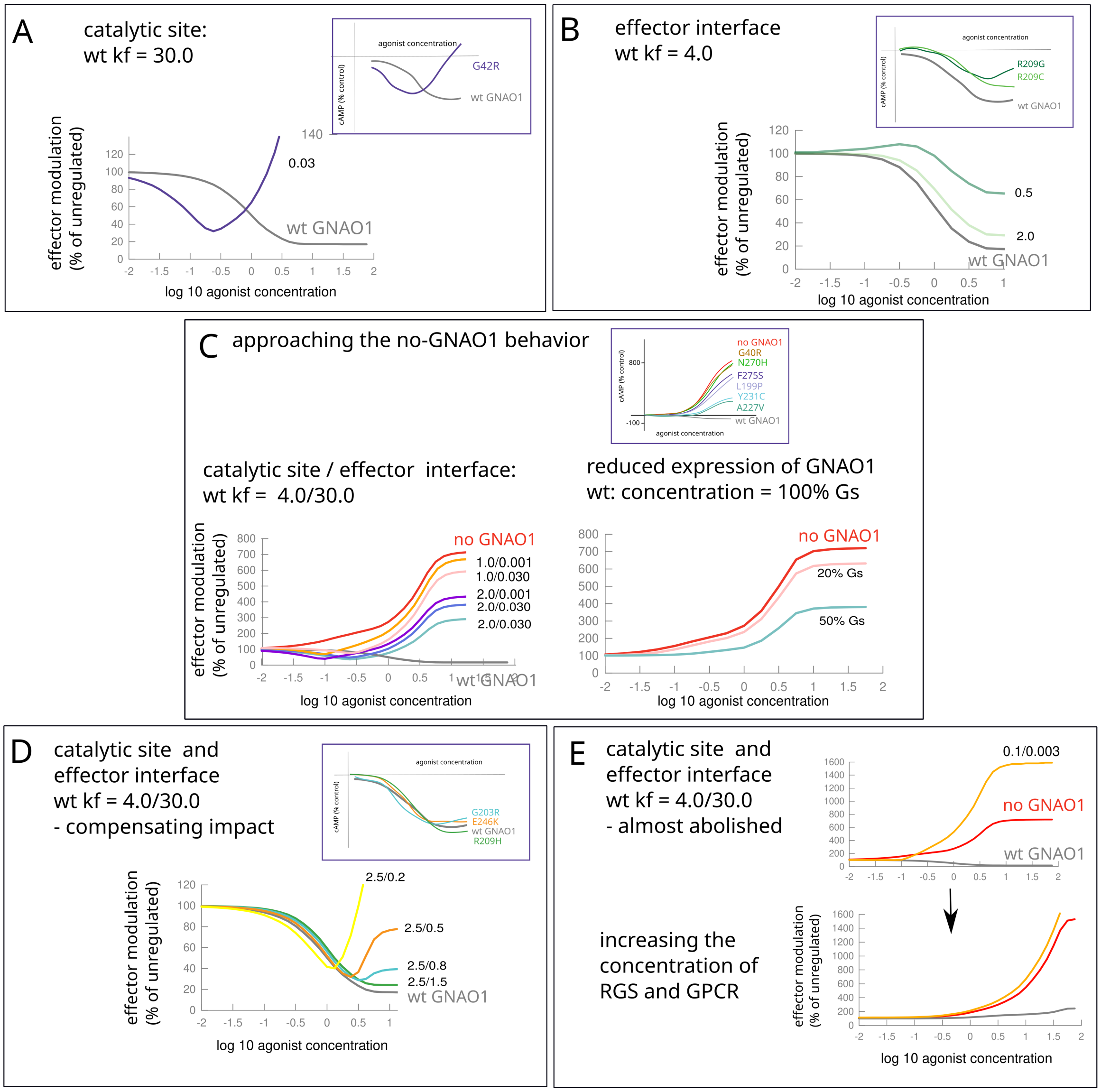


**Fig. S1. Reproducing mutant response to agonist concentration, as reported by Feng *et al.*, (18)**, Fig. 2. The numbers next to each curve show the modified forward constant, for which the wild-type value is given in the panel title. Insets: outline of results from Feng *et al.*

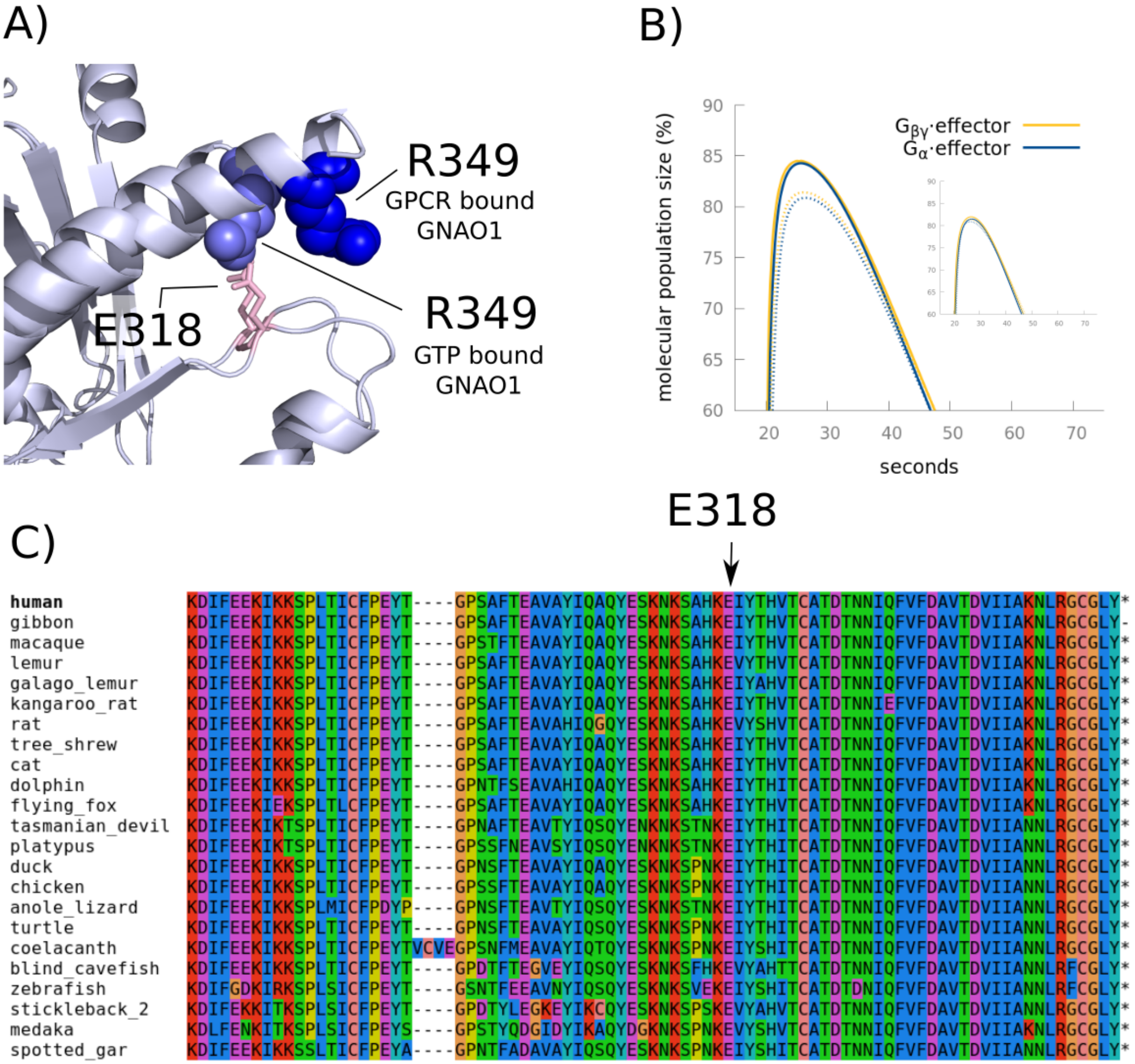


**Fig. S2. Variants with ambiguous phenotype, example: R349-E318 as a latch for non-interacting N-terminal helix.** A) R349 (light blue) forms a salt bridge with E318 (pink) in a nucleotide bound state (when not reacting with GPCR; after structure with PDB identifier 1azs). In a GPCR bound state (GPCR not shown) R349 is facing away from E318 and into the solvent (PDB:3sn6). B) Main figure shows the wild type (dotted) and mutant signal (full line) that increases the rate of GTP to GDP exchange. Inset: rescuing the signal by lowering the amount of agonist. Even if the GTP to GDP exchange was 200 times faster as a consequence of R349Q mutation, the increase in the signal would still be minor, and easily recovered by a modest reduction in the agonist concentration. C) E318 is conserved in all vertebrate homologues of GNAO1.

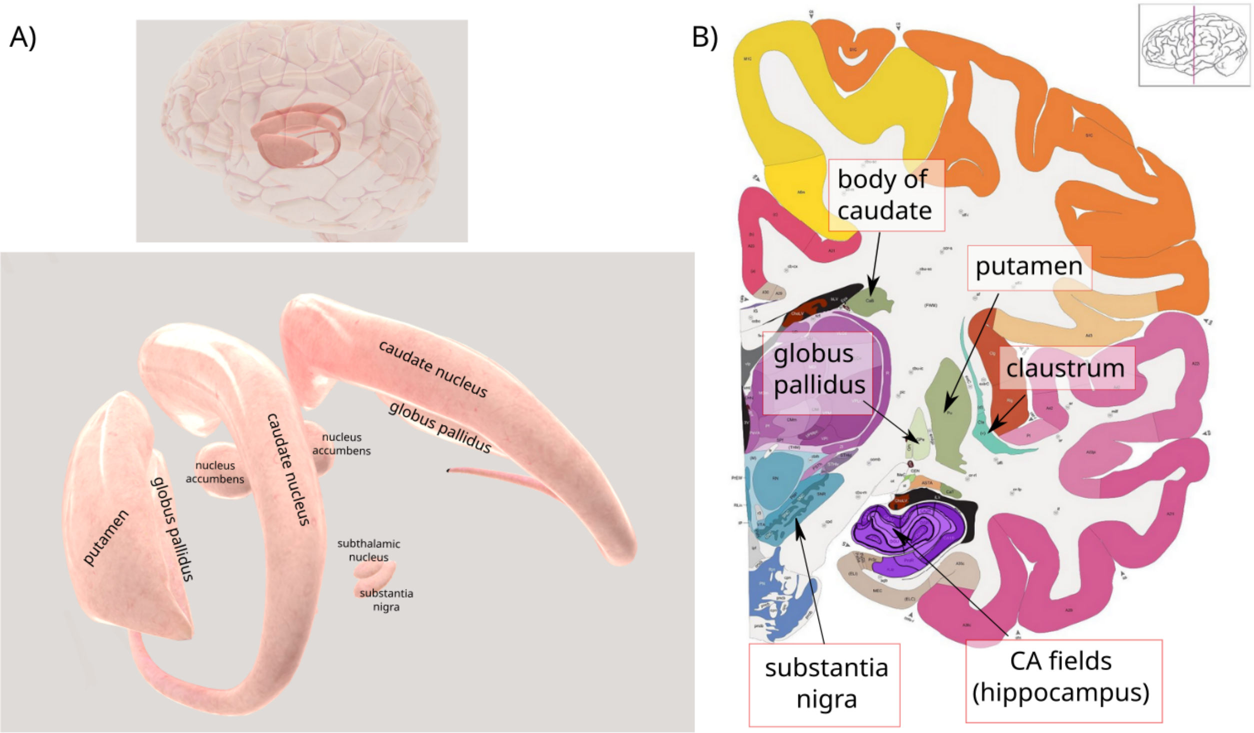


**Fig. S3. Regions of the human brain expressing GNAO1.** A) The topology of basal ganglia, a group of brain nuclei responsible for the modulation of the beginning and end of movement sequences (22). The caudate nucleus and putamen are collectively referred to as the striatum. Caudate and putamen, in particular, are the input region of the basal ganglia (22), and the primary place of expression of GNAO1 (Table S3). Image © Society for Neuroscience (2017), available from brainfacts.org; annotation added for this publication. B) Nomenclature used for the top expressing locations in Tables S3, S4, S1, S2. Image: © 2010 Allen Institute for Brain Science. Allen Human Brain Atlas (23), available from: human.brain-map.org, adult human anatomical reference atlas; annotation added for the purposes this publication.

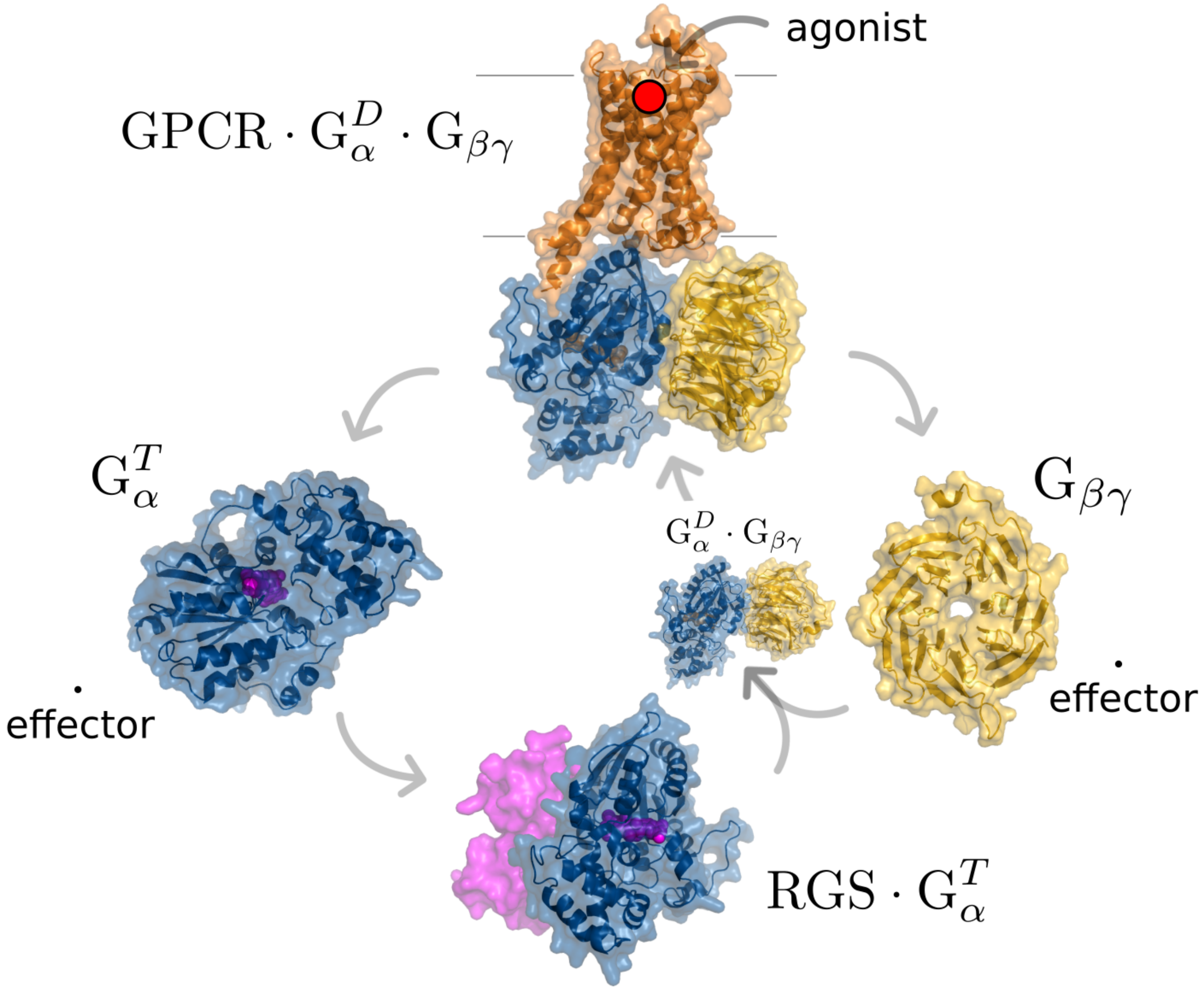


**Fig. S4. Schematic representation of the activated GPCR system.** See Introduction in the main text for the molecular name acronyms. Dots indicate a molecular complex. Superscript $D/T$: bound GDP/GTP. GPCR activated by its agonist, shown as red dot, functions as a guanine exchange factor (GEF) for G$\alpha$. After G-trimer binds to activated GPCR, GDP is released from G$\alpha$, and G trimer dissociates - both from GPCR and internally into activated, GTP bound, G$\alpha$ (blue; GTP indicated in pink) and G$\beta\gamma$ (yellow). Both units immediately associate with their effectors (downstream signaling partners). GTP-bound G$\alpha$ dissociates with some small probability from its effector, which allows it to interact with RGS. RGS speeds up the GTP hydrolysis, and now GDP-bound $G_\alpha$ becomes the preferred interaction partner for G$\beta\gamma$, and the G trimer re-forms. The fraction of time that G$\alpha$ and G$\beta\gamma$ spend interacting with their effectors depends on the ratio of association/dissociation rates for each interaction. The same is true for the time it takes for the system to return to its original state, once the agonist is removed.

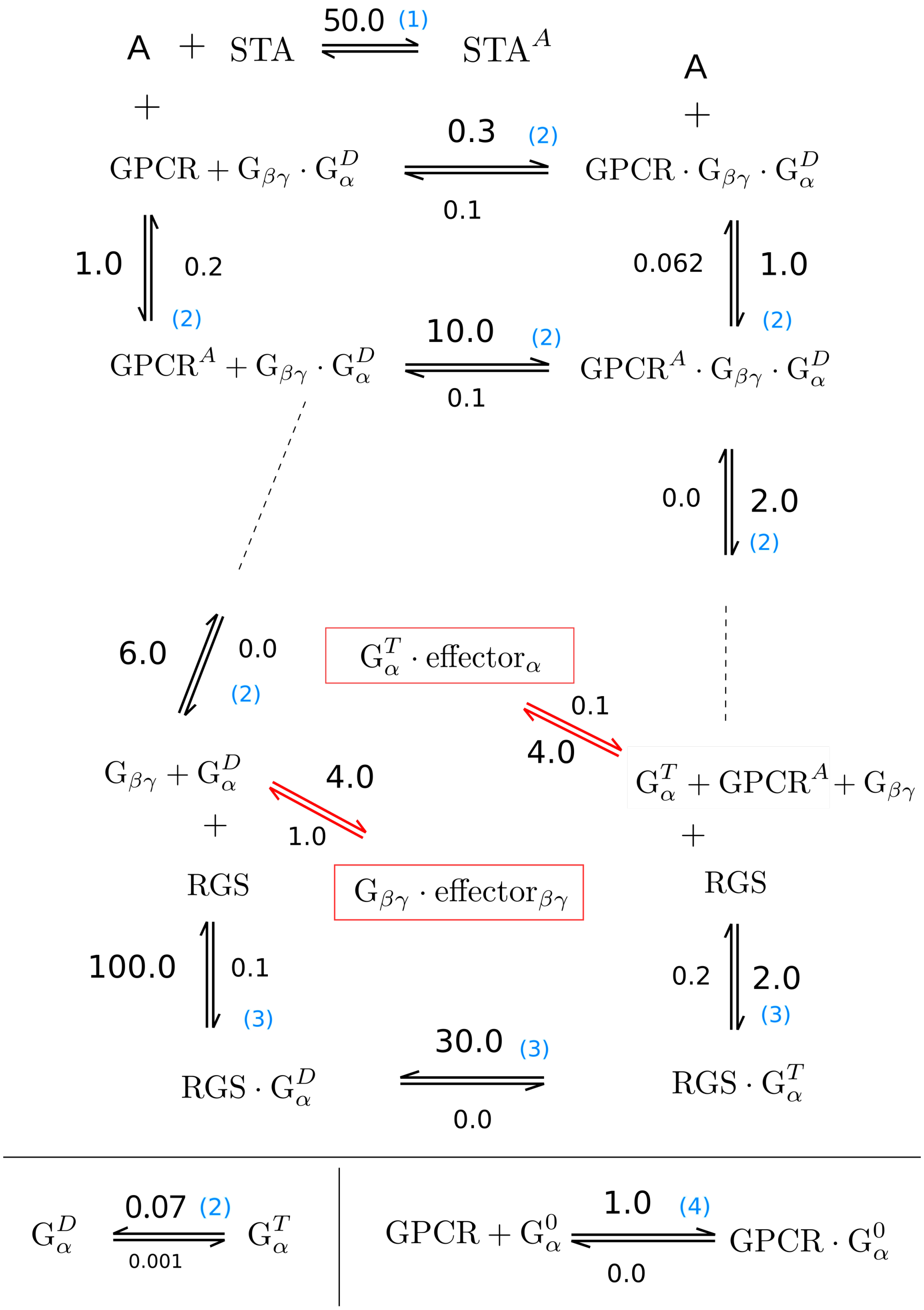


**Fig. S5. GPCR reaction cycle - parametrization for the wild type system.** Disease-related mutations modify these numbers. In the main text, Figs. 4-6 we use modified values to learn how the system reacts to the mutation-induced changes. G: G-protein. GPCR: G-protein coupled receptor. RGS: regulator of G -signaling STA: signal terminating agent. Superscript indicates binding of small ligands: $D$ stands for GDP, $T$ for GTP, and $A$ for generic agonist. Subscripts: subunits of G-protein trimer. Units for the indicated reaction rates: $s^{-1}$ for the first order, and $(\mu Ms)^{-1}$ for the second order reactions. The numbers in the brackets refer to the sources of parameter values: **Source 1:** STA can stand for either acetylcholinesterase in the case of acetylcholine signaling, or GAT proteins in the case of GABAergic signaling. In either case, the reaction rates were set arbitrarily, in order to model the narrow input signal. For acetylcholinesterase activity see (24). In all cases Mg+ ions are assumed to be present and bound to the G$\alpha$ in its catalytic pocket. **Source 2:** Parameters for biding of the agonist and G$\alpha$ to GPCR, for G-trimer formation, and catalysis rate with and without RGS from (14). **Source 3:** Parameters for dissociation of receptor from $G_\alpha^T$ complex, for RGS binding, and for GTP hydrolysis from (13). **Source 4:** Parametrization for "empty pocket" mutant, (15).

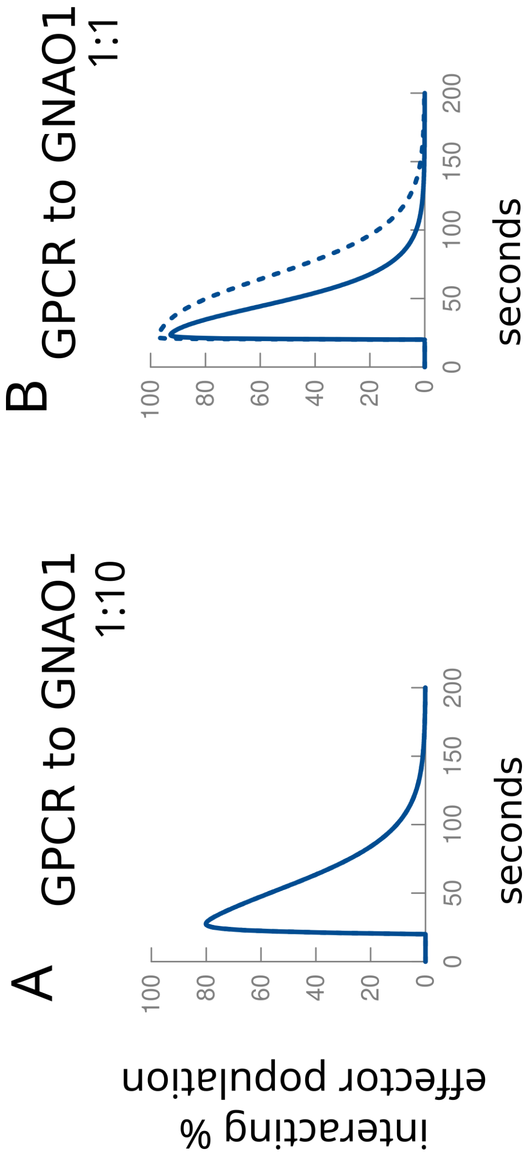


**Fig. S6. The effect of null-mutants.** The mutations that eliminate the mutants completely from the pool of available GNAO1 might either have no effect (A) or result in a weakened concentration (B). **A)** GNAO1 concentration that saturates GPCR. **B**) Comparable GNAO1 and GPCR concentrations. Dashed: wildtype (equals to the case when the mutant is present in A). By comparison of the healthy and affected populations, we surmise that the former might be the case (see the main text, "Null mutants"). The behavior of G$\beta\gamma$ would also depend on the relative ratio of the to G$\alpha$. See S7 below.

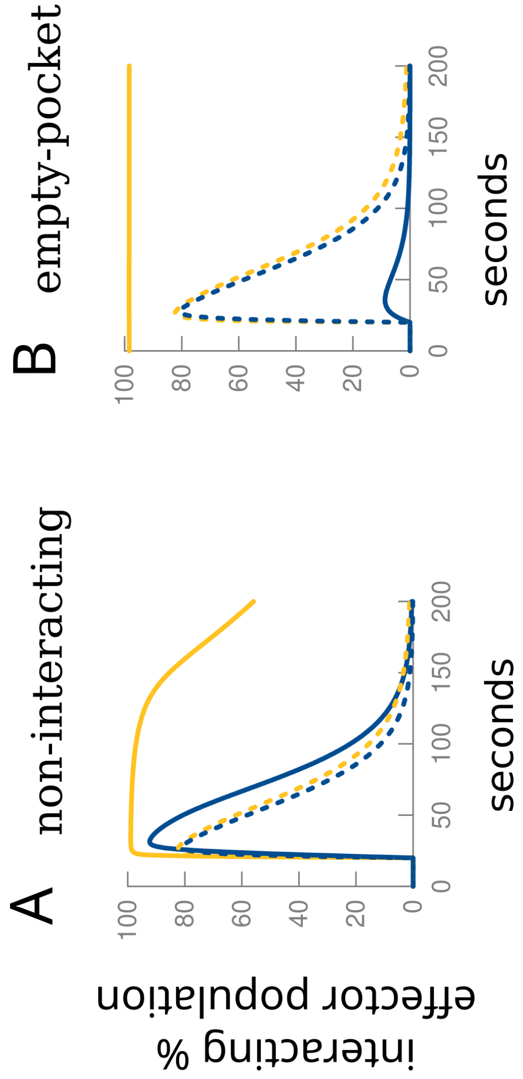


**Fig. S7. Exotic mutants and their effects.** A) Non-interactor. B) Empty pocket.

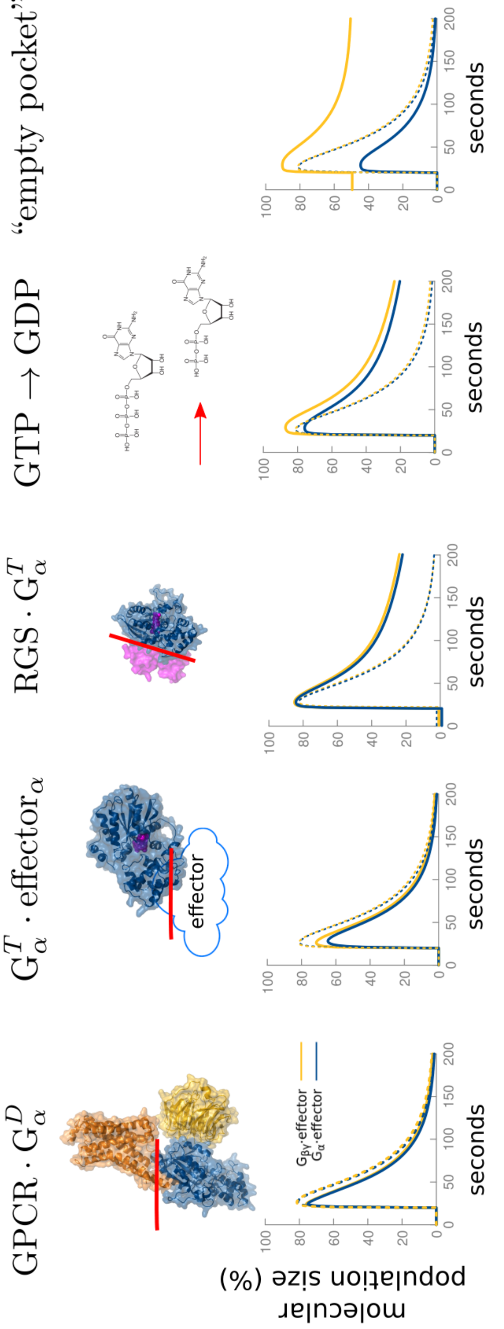


**Fig. S8. The main features of the simulation are not strongly sensitive on the ratio of GPCR:G protein.** Making GPCR's 10-fold more abundant than in the simulations shown in the main text (Figs. 4 and 6) does not change the main features of the signal.

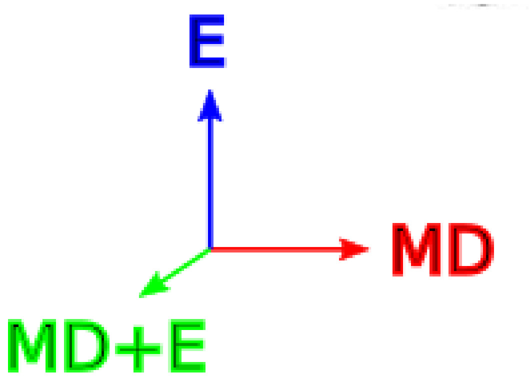


**Fig. S9. RGB phenotype labeling scheme used in the main text.** Used to indicate that for some positions different phenotypes have been reported in literature. For each position three values were calculated: $MD$ = number of patients with MD-only phenotype, $E$ = number of patients with E-only phenotype, and $C$ = number of patients with combined phenotype. With $N = \sqrt{MD^2 + E^2 + C^2}$, the color was assigned according to RGB coloring scheme with RGB vector evaluated as $(MD/N, C/N, E/N)$, thus making pure MD completely red, pure E completely blue, and pure MD+E completely green, with all cases of mixed reports having color somewhere in between. For example, the position for which one patient was reported as E, and one as E+MD would be colored cyan (blue + green).

 Ivana Mihalek, Jeff L. Waugh, Meredith Park, Saima Kayani, Annapurna Poduri, Olaf Bodamer

**Table S1. Coexpression fo GNAO1 and RGS according to The Allen Human Brain Atlas (23) data pre-processed by (25). The numbers shown are negative log p-values for the expression in particular brain region (25). Only the members of RGS family for which data exist are shown. For the nomenclature of the top expressing regions see Fig. S3**

| region | GNAO1 | RGS2 | RGS6 | RGS8 | RGS9 | RGS11 | RGS14 | RGS19 | RGS20 |
|---|---|---|---|---|---|---|---|---|---|
| head of caudate nucleus, right | 1.63 | 1.53 | -1.25 | 1.08 | 1.66 | -1.27 | 1.53 | -1.51 | 2.16 |
| head of caudate nucleus, right | 1.63 | 1.53 | -1.25 | 1.08 | 1.66 | -1.27 | 1.53 | -1.51 | 2.16 |
| head of caudate nucleus, right | 1.63 | 1.53 | -1.25 | 1.08 | 1.66 | -1.27 | 1.53 | -1.51 | 2.16 |
| body of caudate nucleus, right | 1.47 | 0.92 | -1.62 | 0.91 | 2.03 | -0.86 | 1.82 | | 1.66 |
| body of caudate nucleus, right | 1.47 | 0.92 | -1.62 | 0.91 | 2.03 | -0.86 | 1.82 | | 1.66 |
| body of caudate nucleus, right | 1.47 | 0.92 | -1.62 | 0.91 | 2.03 | -0.86 | 1.82 | | 1.66 |
| tail of caudate nucleus, right | 1.37 | | -0.86 | 0.95 | 1.54 | -0.93 | 1.41 | | 1.57 |
| tail of caudate nucleus, right | 1.37 | | -0.86 | 0.95 | 1.54 | -0.93 | 1.41 | | 1.57 |
| tail of caudate nucleus, right | 1.37 | | -0.86 | 0.95 | 1.54 | -0.93 | 1.41 | | 1.57 |
| substantia nigra, pars compacta, right | 1.28 | | 0.83 | | | | | | |
| putamen, left | 1.19 | 1.21 | -1.53 | 0.96 | 1.96 | | 1.58 | -1.32 | 1.52 |
| tail of caudate nucleus, left | 0.96 | 1.20 | -1.23 | 1.11 | 2.13 | -1.03 | 1.87 | -1.02 | 1.52 |
| tail of caudate nucleus, left | 0.96 | 1.20 | -1.23 | 1.11 | 2.13 | -1.03 | 1.87 | -1.02 | 1.52 |
| tail of caudate nucleus, left | 0.96 | 1.20 | -1.23 | 1.11 | 2.13 | -1.03 | 1.87 | -1.02 | 1.52 |
| substantia nigra, pars reticulata, right | 0.90 | | | | | | | 1.70 | |

**Table S2. Coexpression fo GNAO1 and RGS according to The Allen Mouse Brain Atlas (23), in the mouse brain regions corresponding to basal ganglia in human. Data pre-processed by (25). The numbers shown are negative log p-values for the expression in particular brain region (25). Only the same member members of RGS family as in Table S1 are shown.**

| region | GNAO1 | RGS2 | RGS6 | RGS8 | RGS9 | RGS11 | RGS14 | RGS19 | RGS20 |
|---|---|---|---|---|---|---|---|---|---|
| laterostriatal stripe | 3.19 | 1.15 | | 1.09 | 1.56 | 1.30 | | -1.29 | 1.47 |
| intermediate stratum of SeStr | 1.63 | 1.14 | | | 1.68 | | | | |
| accumbens nucleus, shell domain | 1.62 | 1.14 | | | 1.68 | | | | |
| putamen | 1.18 | 1.94 | | 1.04 | 1.75 | 1.16 | | | 1.98 |
| intermediate stratum of Str | 1.08 | 1.85 | | 1.01 | 1.71 | 1.09 | | | 1.90 |
| plexiform layer of TuStr | 1.06 | | | 1.56 | 2.03 | | | | 1.26 |
| striatum (corpus striatum) | 1.01 | 1.75 | | | 1.72 | | | | 1.90 |
| mantle zone of Str | 1.01 | 1.75 | | | 1.72 | | | | 1.90 |
| nucleus of the inferior collicular brachium, caudal part | -1.10 | | | | | | | | -1.16 |
| Spinal nucleus of the trigeminal, caudal part | -1.25 | | | | | | | 1.01 | |
| r10 part of spinal trigeminal nucleus, caudal part | -1.38 | | | | | | | 1.06 | |
| r11 part of spinal trigeminal nucleus, caudal part | -1.77 | | | | | | | | |

**Table S3. Coexpression of GNAO1 and adenylyl cyclase genes according to The Allen Human Brain Atlas (23) data pre-processed by (25). The numbers shown are negative log p-values for the expression in particular brain region (25). Data from ADCY2 and ADCY10 not shown: ADCY10 has no data available, and ADCY2 appears only weakly anticorrelated at a few points. CA fields refer to hippocampal sub-regions (23). For the nomenclature of the top expressing regions see Fig. S3**

| region | GNAO1 | ADCY1 | ADCY3 | ADCY4 | ADCY5 | ADCY6 | ADCY7 | ADCY8 | ADCY9 |
|---|---|---|---|---|---|---|---|---|---|
| claustrum, right | 2.62 | | | | | | 1.42 | | |
| claustrum, left | 2.38 | | | -0.83 | | | 1.23 | | |
| CA2 field, right | 1.84 | | | -0.89 | | | | | 1.63 |
| putamen, right | 1.81 | | 1.24 | | 2.38 | | | -1.46 | 0.98 |
| head of caudate nucleus, right | 1.63 | | 1.65 | | 2.01 | | | -1.67 | 1.18 |
| medial habenular nucleus, left | 1.56 | -2.50 | -3.31 | | -2.34 | | | | |
| CA3 field, right | 1.56 | | | | | | | | 1.85 |
| body of caudate nucleus, right | 1.47 | | 1.76 | | 2.17 | | | -1.29 | 0.88 |
| tail of caudate nucleus, right | 1.37 | | 1.27 | | 1.49 | | | -1.07 | 1.13 |
| substantia nigra, pars compacta, right | 1.28 | -0.93 | | | | | 1.02 | | |
| subcuneiform nucleus, right | 1.27 | | | | | | | | |
| rostral group of intralaminar nuclei, right | 1.26 | | 1.35 | | | | | | |
| lateral habenular nucleus, left | 1.20 | | | | | -0.91 | | | |
| CA4 field, right | 1.19 | | | | | | | | 1.42 |
| putamen, left | 1.19 | | 1.44 | | 1.81 | | | -1.29 | 0.99 |
| Edinger-Westphal nucleus, right | 1.15 | -1.29 | | -1.36 | | | | -1.35 | |
| basolateral nucleus, left | 1.13 | | | | | | | | |
| CA3 field, left | 1.09 | | | | | | | | 1.94 |
| medial group of nuclei, right | 1.08 | | | | | | | 0.95 | |
| CA1 field, right | 1.04 | | | | | | | | 0.89 |
| basomedial nucleus, left | 1.03 | | | | | | | | |
| lateral nucleus, right | 1.01 | | | | | | 0.99 | | |
| lateral hypothalamic area, mammillary region, right | 0.96 | | | | | | | | -1.32 |
| tail of caudate nucleus, left | 0.96 | | 1.54 | | 1.67 | | | -1.46 | 1.39 |
| superior colliculus, right | 0.94 | | -1.16 | | | -1.51 | | | |
| zona incerta, right | 0.93 | | | | | | | | -1.45 |
| oculomotor nuclear complex, right | 0.91 | | -1.43 | | | -1.50 | | | 1.02 |
| substantia nigra, pars reticulata, right | 0.90 | | -1.05 | | | -1.09 | | | |
| anterior group of nuclei, right | 0.87 | | | | | -1.35 | | 1.10 | |
| paraterminal gyrus, right | 0.85 | | | | | | | -1.30 | |
| amygdalohippocampal transition zone, right | 0.83 | | | | | | | | |
| pontine reticular formation, right | 0.83 | | -0.86 | | | -1.08 | | | |
| CA4 field, left | 0.83 | | | | | | | | 1.53 |

**Table S4. Coexpression of GNAO1 and adenylyl cyclase genes in the mouse brain regions corresponding to basal ganglia in human. From The Allen Mouse Brain Atlas (26) data pre-processed by (25).**

| region | GNAO1 | ADCY1 | ADCY3 | ADCY4 | ADCY5 | ADCY6 | ADCY7 | ADCY8 | ADCY9 |
|---|---|---|---|---|---|---|---|---|---|
| laterostriatal stripe | 3.19 | | | | 1.64 | | | -1.58 | |
| intermediate stratum of SeStr | 1.63 | | | | 1.41 | | | | |
| accumbens nucleus, shell domain | 1.62 | | | | 1.41 | | | | |
| putamen | 1.18 | | | | 2.24 | | | -1.57 | |
| intermediate stratum of Str | 1.08 | | | | 2.05 | | | -1.36 | |
| plexiform layer of TuStr | 1.06 | | 1.19 | 2.59 | 2.05 | | | | |
| striatum (corpus striatum) | 1.01 | | | | 2.01 | | | -1.44 | |
| mantle zone of Str | 1.01 | | | | 2.01 | | | -1.45 | |

**Table S5. Default concentrations used in GPCR cycle calculation. For the agonist and the signal terminating agent the time when they appear in the simulation is also indicated. Signal terminating agent (STA) is a generic device to return the system to its equilibrium distribution of molecular species. The agonist/STA pair can be, for example, acetylcholine and acetylcholinesterase.**

| Component | Concentration ($\mu$M) |
|---|---:|
| $G_\alpha$, wild-type | 25 |
| $G_\alpha$, mutant | 25 |
| GPCR | 5 |
| G$\beta\gamma$ | 50 |
| RGS | 30 |
| $G_\alpha$ effector | 50 |
| G$\beta\gamma$ effector | 50 |
| agonist | 60 @ 20.0s |
| signal terminating agent | 120 @ 20.1s |

**Movie S1. The Supplementary Movie 1 contains the visualization of GNAO1 protein structure, its multiple interaction interfaces, and the location of disease-causing mutations relative to them.**

**SI Dataset S1 (supplementary_table_1.xlsx)**

The Supplementary Table 1 contains the list of all disease variants we have collected, their impact on the protein level and the reaction to therapy, cross-referenced with PubMed and DrugBank sources. The legend can be found in the second sheet of this document.
